# Adolescent Social Isolation Reprograms the Medial Amygdala: Transcriptome and Sex Differences in Reward

**DOI:** 10.1101/2020.02.18.955187

**Authors:** Deena M. Walker, Xianxiao Zhou, Aarthi Ramakrishnan, Hannah M. Cates, Ashley M. Cunningham, Catherine J. Peña, Rosemary C. Bagot, Orna Issler, Yentl Van der Zee, Andrew P. Lipschultz, Arthur Godino, Caleb J. Browne, Georgia E. Hodes, Eric M. Parise, Angélica Torres-Berrio, Pamela J. Kennedy, Li Shen, Bin Zhang, Eric J. Nestler

**Affiliations:** Nash Family Department of Neuroscience and Friedman Brain Institute, Icahn School of Medicine at Mount Sinai, New York, NY 10029; Department of Genetics and Genomic Sciences, Icahn School of Medicine at Mount Sinai, New York, NY 10029; Department of Psychology, The University of California Los Angeles, Los Angeles CA, 90095

## Abstract

Adolescence is a sensitive window for reward- and stress-associated behavior. Although stress during this period causes long-term changes in behavior in males, how females respond is relatively unknown. Here we show that social isolation stress in adolescence, but not adulthood, induces persistent but opposite effects on anxiety- and cocaine-related behaviors in male vs. female mice, and that these effects are reflected in transcriptional profiles within the adult medial amygdala (meA). By integrating differential gene expression with co-expression network analyses, we identified crystallin mu (Crym), a thyroid-binding protein, as a key driver of these transcriptional profiles. Manipulation of *Crym* specifically within *adult* meA neurons recapitulates the behavioral and transcriptional effects of social isolation and re-opens a window of plasticity that is otherwise closed. Our results establish that meA is essential for sex-specific responses to stressful and rewarding stimuli through transcriptional programming that occurs during adolescence.

## INTRODUCTION

In many mammals, sex differences in behavior are required for the perpetuation of the species and are expressed as differences in mating and reproductive strategies. Individual variations in reproductive strategies are known to be dependent on environmental factors and rely on programming mediated by experience-dependent plasticity (Bergan et al., 2014). Stressful and rewarding stimuli are two key factors that may be especially important for encoding these strategies to optimize sexual behaviors and maximize reproductive success (Walker et al., 2017). However, very little is known about the brain region-specific molecular processes underlying the programming of such strategies.

Adolescence is a sensitive period for programming reward-associated behaviors particularly in males (Sisk, 2016; Walker et al., 2019) and a time of heightened responsiveness to rewarding and stressful stimuli (Romeo 2013). Recent evidence suggests that the adolescent period may be akin to other critical windows of development, such as sexual differentiation of the brain (Arnold, 2017) and experience-dependent plasticity of the visual system (Hensch, 2005). Stress during adolescence profoundly influences adult behaviors, with adolescent social isolation (SI) in particular altering anxiety- and reward-associated behaviors long-term (Walker et al., 2019): SI increases preference for numerous drugs of abuse including cocaine as well as anxiety- and depression-related behaviors in males (Lukkes et al., 2009; McCormick and Green, 2013; Walker et al., 2019). While decades of research have examined how adolescent SI programs male behavior, much less is known about females (Walker et al., 2019).

Given the adolescent emergence of sex-specific appetitive behaviors, the extended reward circuitry – and the medial amygdala (meA) in particular – is a potential target for the organization that takes place during this time (Walker et al., 2017). First, meA is sexually dimorphic, being larger in males than females (Hines et al., 1992; Meaney et al., 1981), and this dimorphism develops during adolescence (De Lorme et al., 2012). Second, meA is a critical regulator of sex-specific natural reward-associated behaviors (Bergan et al., 2014; Li et al., 2017; Miller et al., 2019; Unger et al., 2015) that emerge during adolescence (Spear, 2000). Third, meA is sensitive to adolescent social stress at the cellular and molecular levels (Cooke et al., 2000; Hodges et al., 2019). Specifically, adolescence SI results in a smaller (more female-typical) meA in male rats compared to group-housed controls (Cooke et al., 2000). Together, these findings suggest that meA may be a key regulator of sex-specific anxiety-like and rewarding behaviors and may be sensitive to perturbations modulated by adolescent experience.

This literature suggests that meA might also be especially sensitive to drugs of abuse, such as cocaine. While there is a body of evidence that the female meA is affected by drugs of abuse in adulthood (Holder et al., 2010; Holder and Mong, 2010; Holder et al., 2015; Williams and Mong, 2017), very little is known about how the male meA responds to drugs of abuse (Knapska et al., 2007). Further, no study has investigated sex-specific transcriptional responses to cocaine in meA in an unbiased manner.

## RESULTS

### Adolescent SI alters sex-specific anxiety- and reward-associated behaviors in adulthood

We first tested the hypothesis that SI exerts long-term effects on anxiety-related behaviors in mice. To determine if any observed effects were specific to adolescence, animals were isolated from P22 – P42 (Figure 1A) or ~P62 – ~P82 (Suppl Figure 1A). Mice were rehoused with their original cage mates at P42 or ~P82, respectively, until behavioral testing occurred at ~P90 (Figure 1A, B) or ~P120 (Suppl Figure 1A, B). Animals were examined for differences in standard anxiety-related behaviors by use of the elevated plus maze

**Figure 1:**
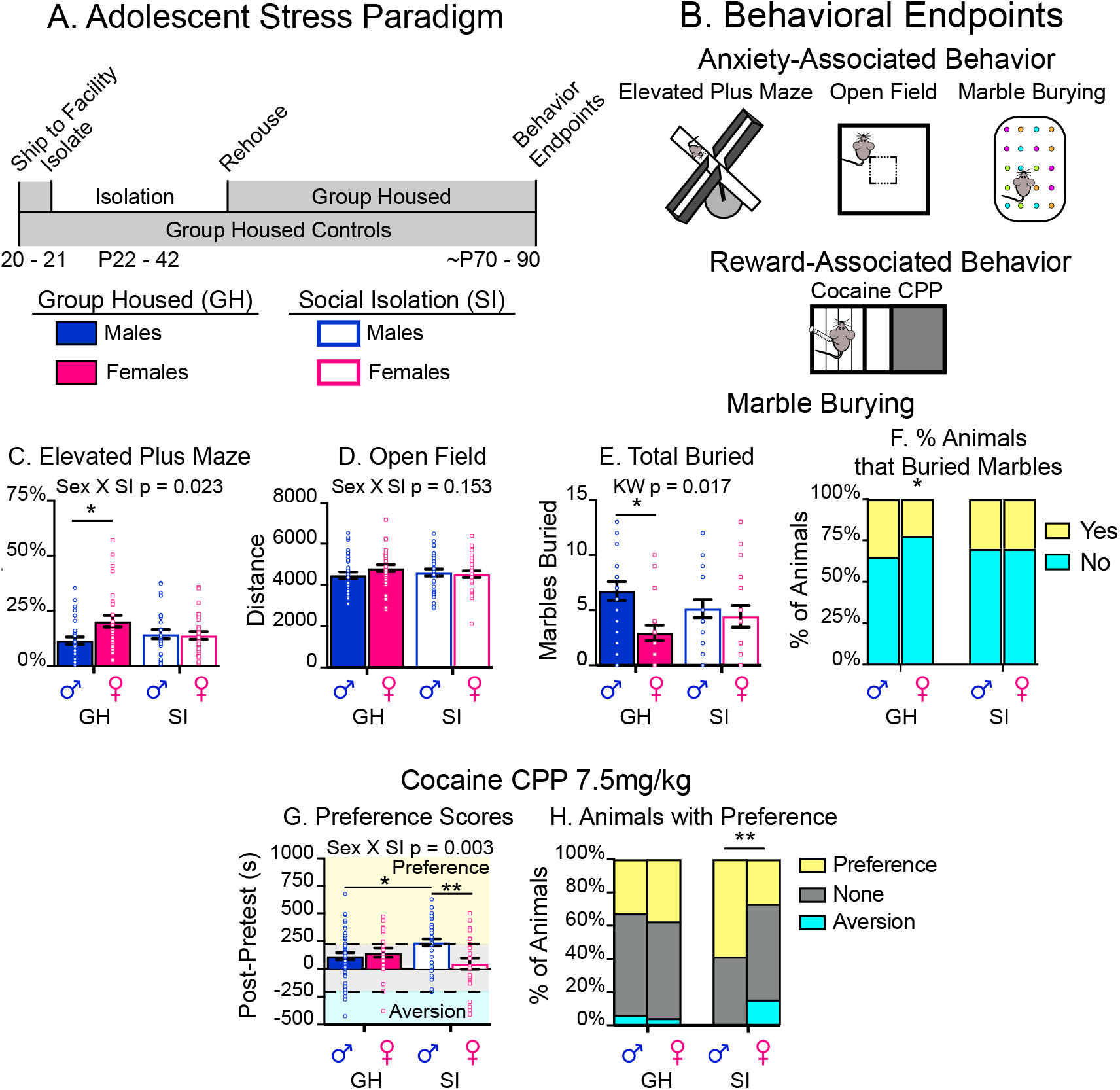
Sexually-Dimorphic Effects of Adolescent SI on Adult Behavior. (A-B) Schematic of experimental design. (C-F) Adolescent SI abolishes known sex differences in elevated plus maze (C) and marble burying (E-F), but has no effect on distance traveled in an open field (D). (G-H) Adolescent SI increases cocaine CPP in males, but decreases it in females (G), resulting in a gain of a sex difference in cocaine-related behavior. The decreased preference for cocaine in SI females is driven by an increase in the number of mice that show an aversion for cocaine (H, blue). Post-hoc significant effects indicated as: *p<0.05; **p<0.001. See also Figure S1.

Sexual differentiation of the brain, and meA in particular, is driven in part by hormone - mediated sex - specific transcriptional programming, which is thought to be permanent after the closure of a critical period of development (McCarthy and Nugent, 2015). There is substantial evidence that sex-specific transcription and behavior are programmed during the post-natal period in numerous brain regions, including meA (Gegenhuber and Tollkuhn, 2019). Evidence also suggests that adolescence may provide a second sensitive window during which sex-specific transcriptional and behavioral programming may occur (De Lorme et al., 2019; Sisk, 2016). However, to our knowledge, further sex-specific transcriptional and behavioral reprogramming has not been observed in adult mammals.

In the present study, we sought to examine whether adolescent experiences, such as SI, can reprogram the meA transcriptome in a sex-specific manner to mediate behavioral changes in a mouse model. Specifically, we answer the following questions: 1. How does adolescent SI alter adult anxiety- and reward-related behavior in males and females? 2. Do transcriptional changes in meA reflect experience-induced differences in behavior? 3. Does targeting a key molecular driver in meA, identified through the integration of several bioinformatic approaches, induce behavioral and transcriptional changes outside the developmental window of sensitivity for sex-specific behaviors.

(EPM), open field (OF) and marble burying tests (Figure 1C-F). SI abolished known sex differences in EPM and marble burying but had no effect on OF (Figure 1C,E,F; EPM: Sex x SI F_(1, 105)_=5.3, p=0.02; Tukey post-hoc: group-housed males [GHM] vs. group-housed females [GHF] p=0.02; total marbles buried: Kruskall-Wallace H_(3, 78)_=10.2, p=0.02; distribution of animals that buried marbles: Chi squared – group-housed males [GHM] (χ^2^=1.8; p=0.2), group-housed females [GHF] (χ^2^ = 5.56; p=0.02), SI males [SIM] (χ^2^=3.2; p=0.07), SI females [SIF] (χ^2^=3.2; p=0.07)). Importantly, SI in adulthood did not affect sex differences in EPM (Sex x SI; F_(1, 64)_ = 0; p=0.98; Sex_;_ F_(1, 64)_ = 31.69; p<0.001). OF (Sex x SI; F (_(1, 63)_ =0.57; p=0.45; Sex_;_ F_(1, 63)_ = 10.44); p<0.001), or marble burying (Sex x SI; F_(1, 64)_=2.31; p=0.13; Sex_;_ F_(1, 64)_=2.96); p=0.09).

While numerous studies have shown that adolescent SI increases preference for drugs of abuse in males, much less is known about females (Walker et al., 2019). We used cocaine conditioned place preference (CPP; 7.5mg/kg) to determine if adolescent SI induces sex-specific effects on reward sensitivity in adulthood (Figure 1G,H). Two-way ANOVA revealed an interaction of sex and SI (Figure 1G; F_(1, 143)_=8.98; p<0.01). Tukey post-hoc analysis indicated that, as predicted, SIM showed increased cocaine CPP over GHM p=0.04), whereas SIF showed a non-significant decrease (GHF vs. SIF: p=0.3) in their preference compared to their GH counterparts.

Adolescent SI thus results in a *gain* of sex difference in cocaine CPP (SIM vs. SIF; p<0.01) and a shift in the proportion of animals that formed a preference for or aversion to cocaine (Figure 1H; GHM vs. GHF χ^2^=0.29; p=0.87; SIM vs. SIF: χ^2^=9.41; p<0.01). In contrast, the same isolation paradigm in adulthood had no effect on cocaine CPP (Sex x SI; F_(1, 37_ =0.09; p=0.77) or the proportion of animals forming a preference in any group (Suppl Figure 1G,H). Together, these results establish adolescence as a sensitive period for developing sexually-dimorphic anxiety- and reward-associated behaviors.

### meA transcriptome after adolescent SI reflects sex-specific behaviors

We hypothesized that meA undergoes lasting sex - specific transcriptional reprogramming in response to adolescent SI that partly underlies the observed behavioral effects. Using RNA-seq, we analyzed transcriptome-wide gene expression changes in meA of GHM/F or SIM/F after acute (1 hr after first dose) or chronic (24 hr after 10^th^ dose) of investigator-administered cocaine (7.5 mg/kg) or saline (Figure 2A,B; Suppl Table 1). We chose these dosing paradigms to determine if SI disrupts sex-specific sensitivity to cocaine (acute) and if long-term cocaine-induced transcriptional alterations (chronic) are disrupted by SI. This approach was based on other studies from our laboratory showing that acute vs. chronic cocaine exposure have different transcriptional effects in other brain regions (Maze et al., 2010; Walker et al., 2018).

**Figure 2:**
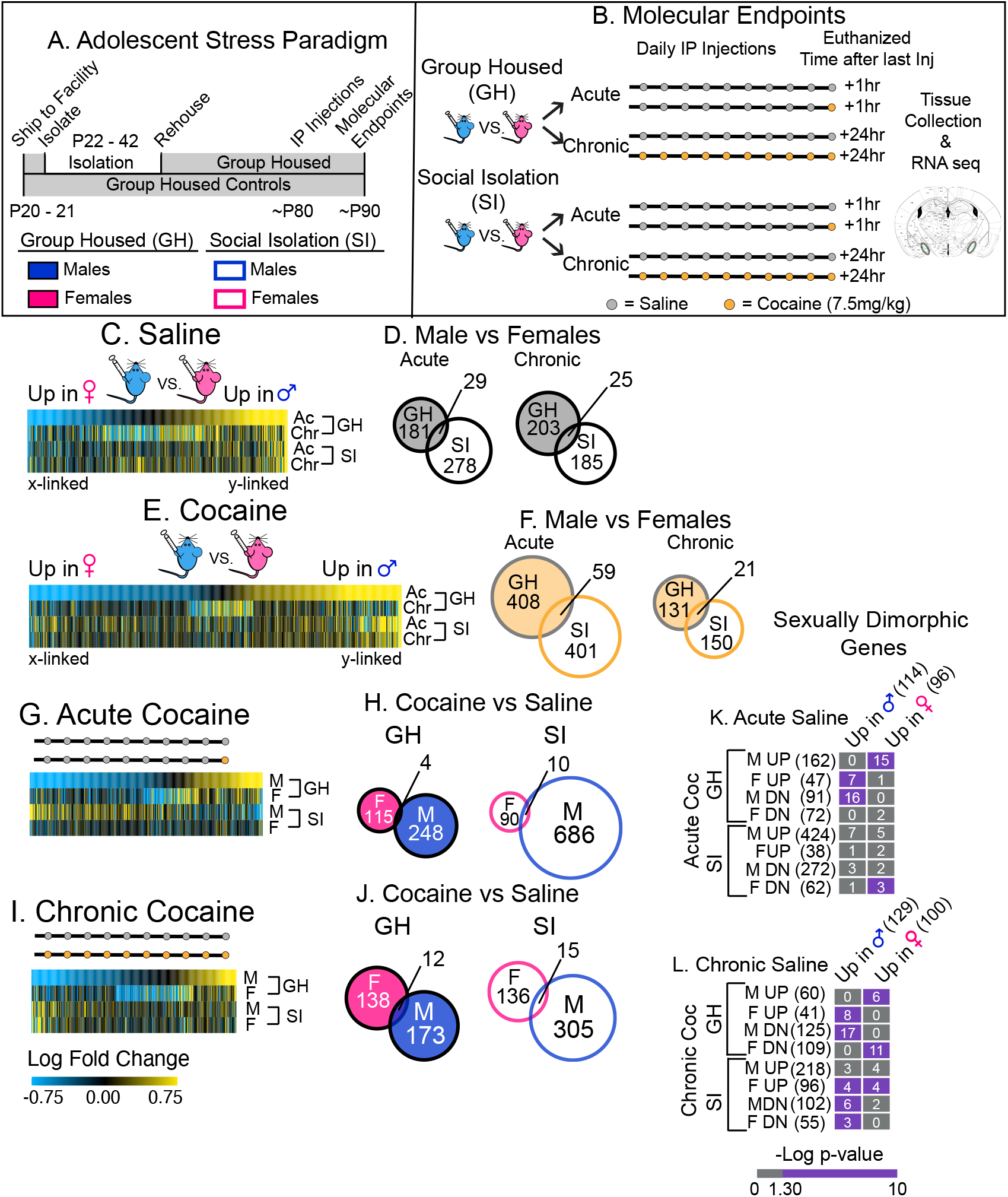
Transcriptional Effects of Adolescent SI in Adult meA Mirror the Behavioral Effects of SI. (A-B) Schematic of experimental design. (C-F) Union heatmaps and Venn diagrams of all sexually-dimorphic DEGs in GHM vs. GHF after acute or chronic saline (C-D) or acute or chronic cocaine (E-F). There are more sex differences in DEGs after acute saline (C-D) or cocaine (E-F) than under chronic conditions. Heatmaps reveal that adolescent SI abolishes sex-specific expression under all conditions. (G-J) Union heatmaps and Venn diagrams of DEGs in GHM/F after acute (G-H) or chronic (I-J) cocaine, each compared to their own saline control. There is very little overlap of DEGs regulated by cocaine in both GH and SI males vs. females. The heatmaps also reveal that adolescent SI induces robust but opposite transcriptional responses to cocaine in meA in males but not females. (K-L) Enrichment of sexually-dimorphic genes in GH animals after acute (K) or chronic (L) saline with those genes altered by cocaine in GH and SI animals. Grids show pairwise comparisons. The color of each box indicates significant enrichment of genes lists; purple = significant enrichment of the two lists (FDR p<0.05). The number in each box indicates the number of overlapping transcripts. Numbers in parentheses = the total number of genes in the list. See also Suppl Table 1 and Suppl Figure 2.

To assess how adolescent SI affected sex differences in meA gene expression, we focused first on genes that are sexually-dimorphic in GH and SI mice after saline (Figure 2C,D and Suppl Figure 2C) or cocaine (Figure 2E,F and Suppl Figure 2D). Union heatmaps and Venn diagrams of differentially-expressed genes (DEGs) between GHM vs. F after acute and chronic saline (Figure 2C,D) or acute and chronic cocaine (Figure 2E,F) reveal that expected sex differences in gene expression (those observed in GH mice) are lost after SI in all treatment groups. Venn diagrams also indicate little overlap of sexually-dimorphic DEGs between GH and SI animals after saline or cocaine (Figure 2D,F). The lack of overlap of DEGs indicates that SI results in both a *loss* (DEGs between GHM vs. GHF are not observed after SI) and *gain* (DEGs between SIM vs. SIF are not observed in GH animals) of sex differences in expression under all conditions. These data show that, parallel to behavioral effects of SI, expected sex differences in gene expression are lost in meA after SI both at baseline and after cocaine.

To examine how meA gene expression in males and females responds to cocaine, we generated union heatmaps of DEGs regulated by cocaine in GHM/F after acute (Figure 2G,H) or chronic (Figure 2I,J) cocaine compared to each group’s saline control (acute or chronic). SIM, but not SIF, displayed an opposite transcriptional response to cocaine compared to GHM and, to a lesser extent, GHF (Figure 2G,I; Suppl Figure 2F). Consistent with prior findings (Barko et al., 2019; Hodes et al., 2015; Labonte et al., 2017; Pena et al., 2019; Seney et al., 2018), Venn diagrams (Figure 2H,J) indicate very little overlap (<10%) of those genes regulated by acute or chronic cocaine in GHM and GHF. Interestingly, SIM but not SIF mount a robust transcriptional response to both acute and chronic cocaine (Figure 2H,J) and union heatmaps of SIM/F response to acute and chronic cocaine (Suppl Figure 2E, F) reveal opposite transcriptional responses in GHM in particular. Together, these data suggest that males and females differ in their meA transcriptional response to cocaine and those differences are lost after SI.

Fisher’s exact tests (FETs) revealed significant enrichment (FDR p<0.05) of sexually-dimorphic genes under saline control conditions in the lists of genes altered by cocaine (acute and chronic) in GH mice (Figure 2K,L). This suggests that cocaine disrupts baseline sex differences in meA gene expression.

### Disruption of sexually-dimorphic gene expression in meA after adolescent SI

The finding that sexually-dimorphic genes in meA are altered by cocaine in GH mice (Figure 2K,L) and are lost after SI (Figure 2C-F) suggests that baseline sex differences in gene expression are disrupted after adolescent SI. We used pattern analysis, an analytical technique employed previously by our laboratory (Walker et al., 2018), to reduce the dimensions of our data and identify genes with patterns of expression that can be categorized based on observed behavior (see Methods). All DEG data were compared to the same baseline (GHF given chronic saline). While many patterns were identified in this dataset (Suppl Table 2), we chose to focus on two patterns: genes that are different between GHM and GHF (Figure 3), and 2) genes that are only regulated in SI mice (Suppl Figure 3). By focusing on these two patterns, we identified genes representing a loss of sex differences (Figure 3A,B) or gain of sex differences (Suppl Figure 3A, B) in response to adolescent SI. We further categorized the sexually-dimorphic gene patterns as follows: “feminized” (Fem) if the SIM expression pattern resembled GHF (Figure 3A); “masculinized” (Mas) if the SIF expression pattern resembled GHM (Figure 2B); “sex difference reversed” (SDR) if the opposite pattern was observed in SIM vs. SIF compared to GHM vs. GHF (Figure 3C); and “sex difference maintained” (SDM) if expression was similar in SI and GH M vs. F (Figure 3D).

**Figure 3:**
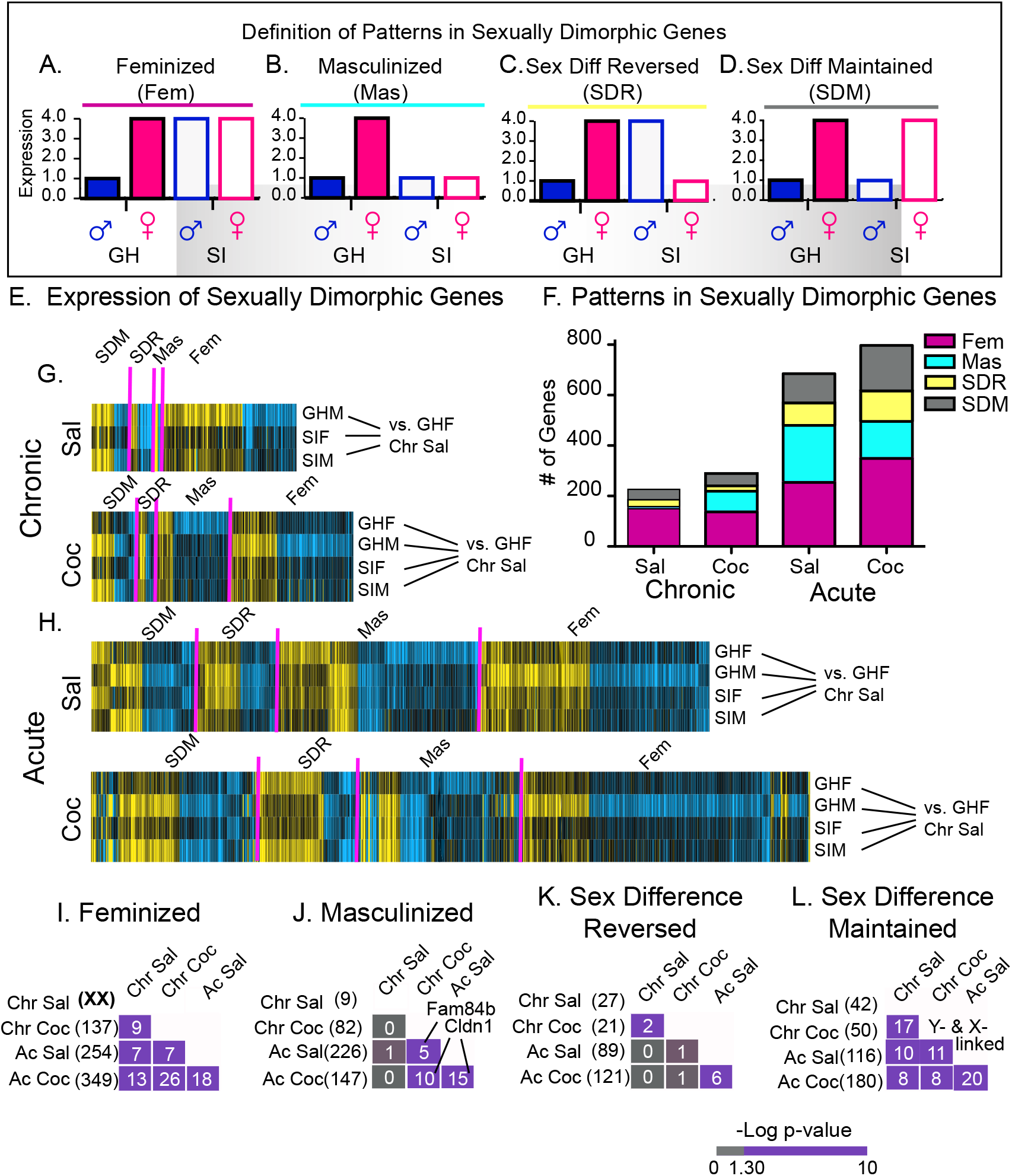
Pattern Analysis Reveals that Sexually-Dimorphic Genes in meA are Feminized by Adolescent SI. (A-D) Theoretical patterns: (A) Feminized (Fem) if SIM expression resembles GHF; (B) Masculinized (Mas) if SIF expression resembles GHM; (C) Sex Difference Reversed (SDR) if a baseline sex difference is reversed by SI; and (D) Sex Difference Maintained (SDM) if a baseline sex difference is maintained after SI. (E-H) Heatmaps of sexually-dimorphic genes that fall into these categories. All comparisons are made to the GHF chronic saline group and expressed as logFC. Pink lines indicate breaks between patterns. Number of sexually-dimorphic genes categorized as each pattern for each treatment paradigm. (I-L) Enrichment of sexually dimorphic genes categorized by their patterns across treatment paradigms. Grids show pairwise comparisons. The color of each box indicates significant enrichment of genes lists; purple = significant enrichment of the two lists (FDR p<0.05). The number in each box indicates the number of overlapping transcripts. Numbers in parentheses = the total number of genes in the list. See also Suppl Figure 3 and Suppl Table 2.

When the data were compared to the same baseline, it is clear that males and females regulate many more transcripts in response to an acute injection of saline or cocaine (Figure 3G,H) compared to chronic conditions (Figure 3E,F), suggesting that gene transcription in males and females responds robustly in meA to handling and IP injections, even after 10 days of habituation. The larger number of transcripts affected after acute conditions likely reflects the fact that mice were examined 1 hr after injection in contrast to 24 hr used for chronic conditions. This analysis reveals that SI results in a greater proportion of transcripts being feminized in SIM under all four treatment paradigms (Figure 3I; Suppl Figure 3E). These data further support the finding that SI in males results in a more transcriptionally responsive meA and that the loss of sex differences in expression may be driven by transcriptional changes within SIM.

For females, we observed that virtually no transcripts in SIF are masculinized under chronic saline conditions. However, in the other treatment paradigms, ~25% of transcripts are masculinized. This suggests that a second stimulus is necessary to induce male-typical expression of certain transcripts in SIF (Figure 3F,G), which is supported by enrichment analysis of pattern DEGs categorized by their expression profile across treatment paradigms. FETs revealed significant enrichment of transcripts categorized as feminized (Figure 3I), SIM only (Suppl Figure 3G), and SIF only (Figure S3H), but masculinized genes were only enriched after a second stimulus (Figure 3J). These findings indicate that transcripts are programmed to respond to stimuli in a male- or female-typical manner and that SI disrupts this signaling.

### Co-expression analysis reveals that feminization and masculinization of sexually-dimorphic genes disrupts transcriptional networks

Gene co-expression network analysis is a crucial tool for identifying transcripts that are changing together, with such networks providing valuable insight into psychiatric disorders (Bagot et al., 2016; Gaiteri et al., 2014; Labonte et al., 2017; Parikshak et al., 2013). Consequently, to better understand meA transcriptional contributions to sex-specific behaviors, we compared sex-specific co-expression networks between GHM and GHF. We then determined if disruption of sexually-dimorphic gene expression in meA after SI altered gene co-expression on a global scale. Using multiscale embedded gene co-expression network analysis (MEGENA; (Song and Zhang, 2015), we identified gene co-expression networks that correlate with sex-specific effects of SI. Unlike other forms of co-expression analysis, MEGENA not only identifies modules of co-expressed genes but also utilizes a hierarchical model to provide information about co-expression across the entire transcriptome by iteratively identifying clusters of smaller co-expression modules embedded within larger “parent” modules. This hierarchical approach is important, because it provides information about transcriptomic “structure,” including the number of parent modules evident genome-wide.

Modules were constructed for each of the four groups (GHM/F, SIM/F) by collapsing data from all four treatment paradigms (acute/chronic cocaine and saline). We predicted that sex differences in transcription would be reflected in global transcriptomes of GHM and GHF (Figure 3 A,B). In GHM, we identified 10 parent modules (Figure 4A,E,F) comprising fewer genes than their GHF counterparts (Figure 4B,E,F) – corresponding to less global co-expression in GHM vs GHF. Critically, SI dramatically reverses this effect: SIM have fewer parent modules than SIF (Figure 4C-F), and the size of SIM parent modules is closer to GHF (Figure 4F). The most striking effect was in SIF where the number of modules is vastly greater than the other groups and the size of the modules is significantly smaller (p<0.05). These data reveal that adolescent SI disrupts global co-expression in a sex-specific manner.

**Figure 4:**
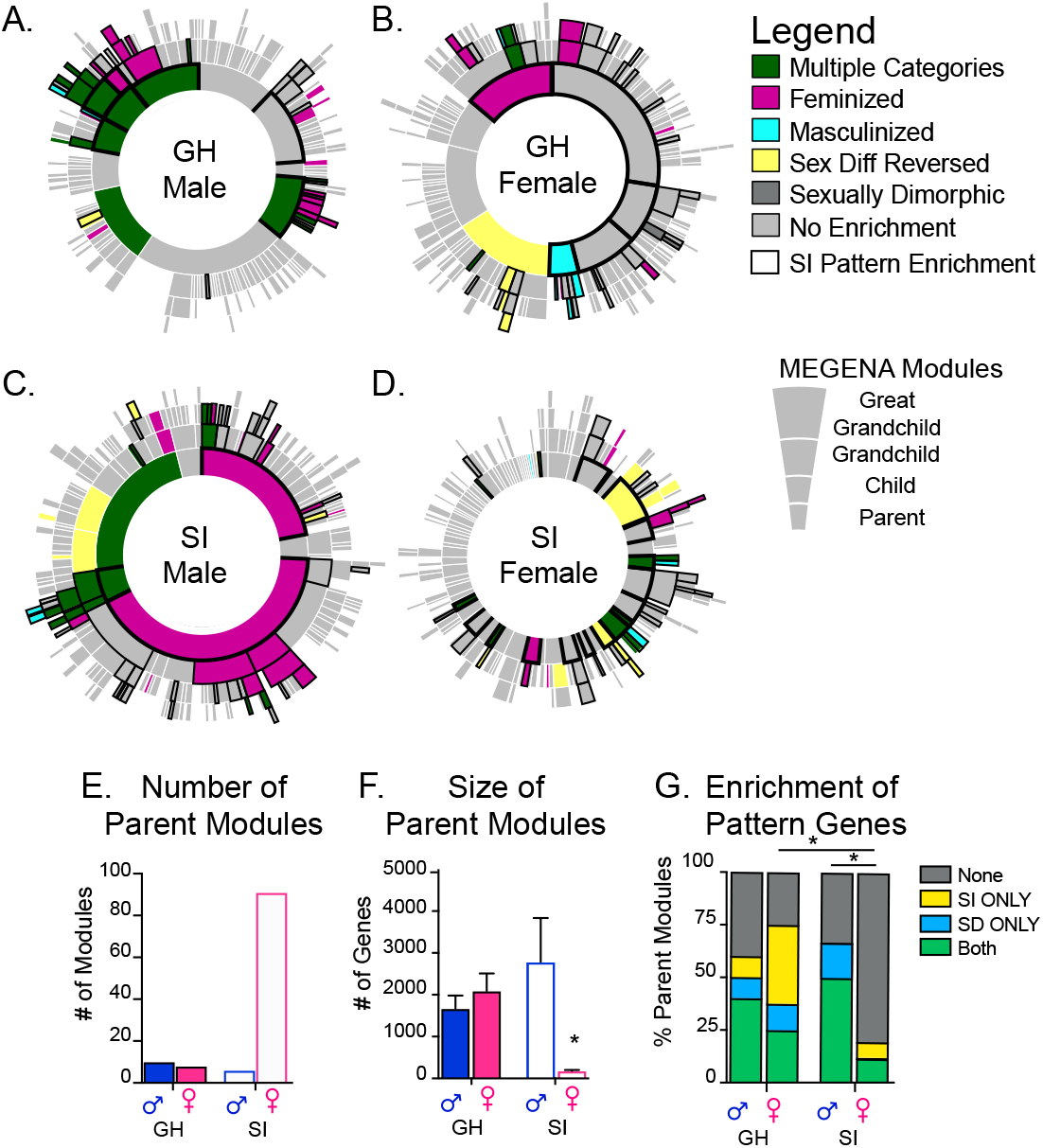
Co-Expression Network Analysis Reveals That Adolescent SI Feminizes the Male Transcriptome in meA. (A-D) Sunburst plots derived from MEGENA of GHM and GHF (B) compared to SIM (C) and SIF (D). The inner, middle and outer rings depict “parent, children, and grandchildren” modules, respectively. Colors indicate enrichment of categories of sexually-dimorphic genes (SD genes) determined from the pattern analysis in each module. Bold outline indicates enrichment of genes only affected in SI animals (SI genes) determined from the pattern analysis. (E-F) Bar graphs of number of parent modules (E) and average number of genes per parent module (F) in each group. (G) Enrichment of SI and SD pattern genes in the parent modules overlaps in males more than females. Percent of modules enriched in sexually-dimorphic (blue), SI-only (Yellow) or both (green) patterns are indicated. *p<0.05; **p<0.001.

To determine if sex-specific disruption of co-expression was driven by a loss or gain of sex differences in DEGs after SI, we determined if modules were enriched in categories of genes identified using patterns analysis (Figure 4A-D,G). We hypothesized that modules enriched in specific categories of sexually-dimorphic genes may reflect modules whose sex-specific co-expression is lost after SI (e.g., enrichment of feminized or masculinized genes; Figure 4G blue) or sex-specific co-expression was gained after SI (enrichment of SI-only genes; Figure 4G yellow). Enrichment of both patterns within a module (Figure 4G, green) would reflect both a gain and loss of sex-specific co-expression and may provide information regarding how SI reprograms sex-specific co-expression of the global transcriptome. Indeed, SIM parent modules (Figure 4C,G) display a high degree of overlap of enrichment of both sexually-dimorphic patterns (mainly feminized) and SI-only patterns, whereas SIF have little enrichment of either pattern (GHF vs. SIF p = 0.028; SIM vs. SIF p=0.032; Figure 4D,G). These strong differences in SIM vs. SIF global co-expression and co-enrichment suggest that SI likely disrupt sex-specific key drivers of transcription in meA.

### Sex-specific key drivers are associated with immune and hormonal signaling

Through an integration of our co-expression, pattern and enrichment analyses, we identified key driver genes that reflect the sex-specific differences in behavior observed after SI. We identified drivers that met the following 5 criteria: 1) conserved in all 4 groups (GHM/F and SIM/F); 2) sex-specific betweenness (a measure of a driver’s location within the module, e.g., in the center or periphery) in GH modules is lost/reduced after SI; 3) sex-specific connectivity in GH modules is lost/reduced after SI; 4) categorized as a sexually-dimorphic or SI-only pattern gene; and 5) the module containing the key driver is enriched in sexually-dimorphic and/or SI-only pattern genes (Figure 5A). We identified 2 key drivers that met all 5 criteria (Figure 5B-I): crystalin mu (*Crym*) and prostaglandin D2 synthase (*Ptgds*). *Crym* is a thyroid hormone-binding protein (Hallen et al., 2011; Vie et al., 1997), a non-canonical transcriptional regulator (Ohkubo et al., 2019), and is neuroprotective in striatum (Francelle et al., 2015), while *Ptgds* is a pro-inflammatory enzyme which catalyzes the production of prostaglandin D (Urade and Hayaishi, 2011). Since we hypothesized that transcriptional changes within meA drive sex-specific changes in behavior, we chose to focus on *Crym* (Figure 5J-M).

**Figure 5:**
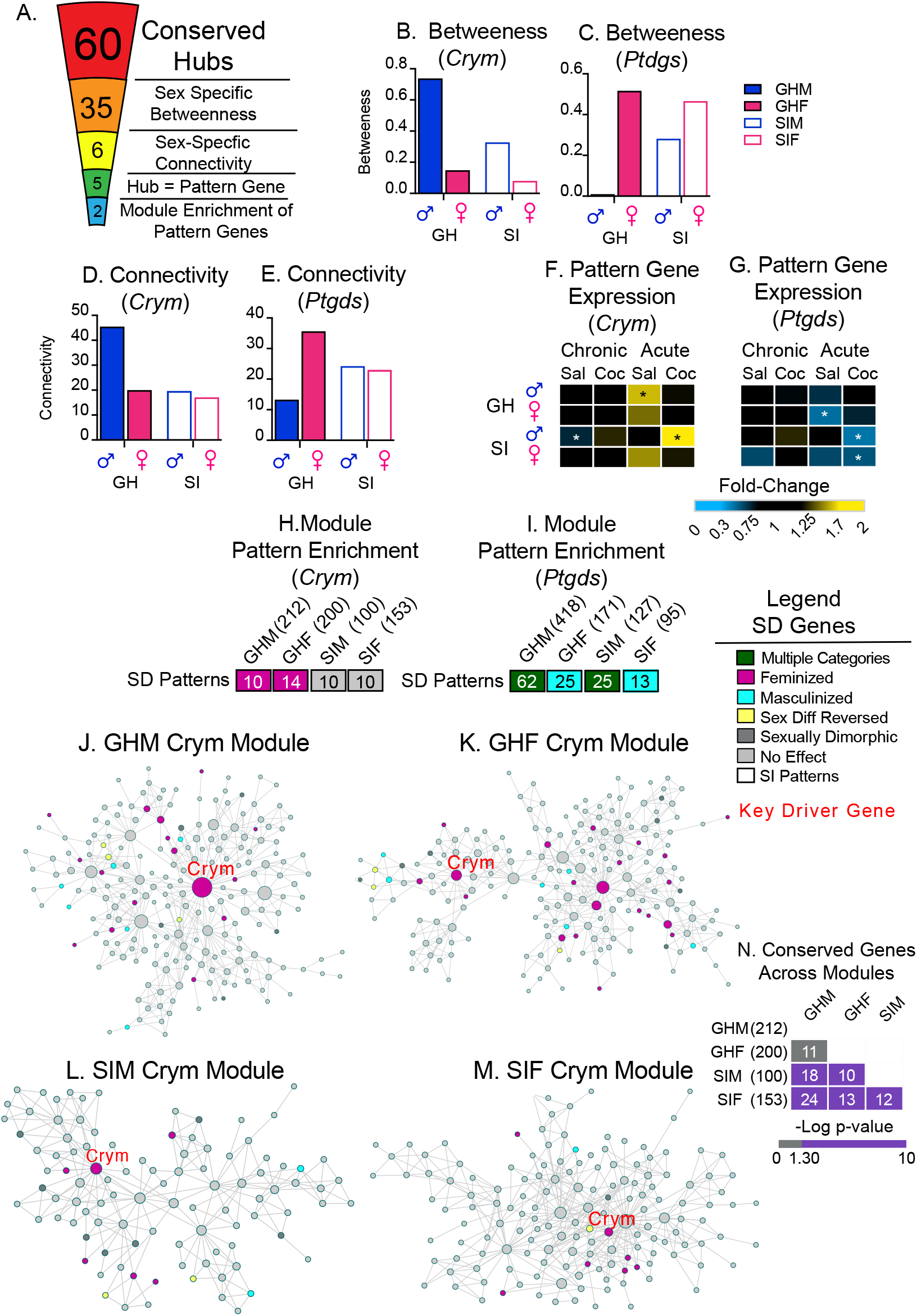
Co-Expression Analysis Reveals *Crym* as a Conserved Key Driver of Sex-Specific Gene Expression. (A) Schematic of criteria used to identify sex-specific key-driver genes. The number in each segment indicates the number of key drivers meeting each criterion. (B-D) *Crym* and *Ptgds* display sex-specific betweenness (B-C), and connectivity (D-E) within their respective modules for GH animals (closed bars). These differences are lost or diminished in modules for SI animals (open bars). (F-G) Fold change in expression compared to same baseline (GHF after chronic saline) from RNA-seq data. *expression differs from baseline. (H-I) Enrichment of sexually-dimorphic and SI-only pattern genes is observed in modules containing *Crym* and *Ptgds,* respectively. Significant enrichment of a category of pattern genes is color coded; gray = no significant enrichment. (J-M) Arachne plots of *Crym* modules in all groups. In each case, *Crym* is a key driver. Sexually dimorphic genes falling into different patterns are color-coded. (N) Enrichment plot of conserved genes across *Crym*-containing modules. Significant enrichment is indicated by purple, number of genes in each module is in parentheses, and number of overlapping genes is in each box. See also Suppl Figure 5.

### *Crym* overexpression (OE) in adult meA recapitulates the effects of adolescent SI

Based on patterns of expression in GH and SI mice (loss of sex difference after acute saline; gain of sex difference after acute cocaine), we hypothesized that *Crym* OE would recapitulate the effects of SI for both anxiety- and cocaine-related behaviors (Figure 5F). AAV vectors (AAV2-hsyn-*Crym*-GFP or GFP alone) were bilaterally injected into meA of adult males and females and behavioral testing was conducted 3 weeks later. *Crym* overexpression was confirmed after behavioral experiments by epifluorescence microscopy and qPCR (Figure 6B). Only those animals that had correct targeting and had at least 1.5 standard deviations greater *Crym* expression than GFP controls were included in the analysis. As predicted, sex differences in EPM (Figure 6C; Sex x SI F_(1,66_=5.11; p=0.029) were observed in GFP controls but not after *Crym* OE in meA (Tukey post hoc: GHF vs GHM p<0.05). Two-way ANOVA revealed no significant effects of *Crym* OE on OF (Sex x SI F_(1,69)_=0.17; p=0.71) or total marbles buried (Sex x SI F_(1,69)_=1.07; p=0.14) but, similar to SI, *Crym* OE increased the proportion of females that buried marbles to male levels (Figure 6F; Chi squared: GFPM (χ^2^=1.2; p=0.27), GFPF (χ^2^=9.97; p<0.01), CrymM (χ^2^ =1.33; p=0.25), and CrymF (χ^2^=0.0; p=1.0).

**Figure 6.**
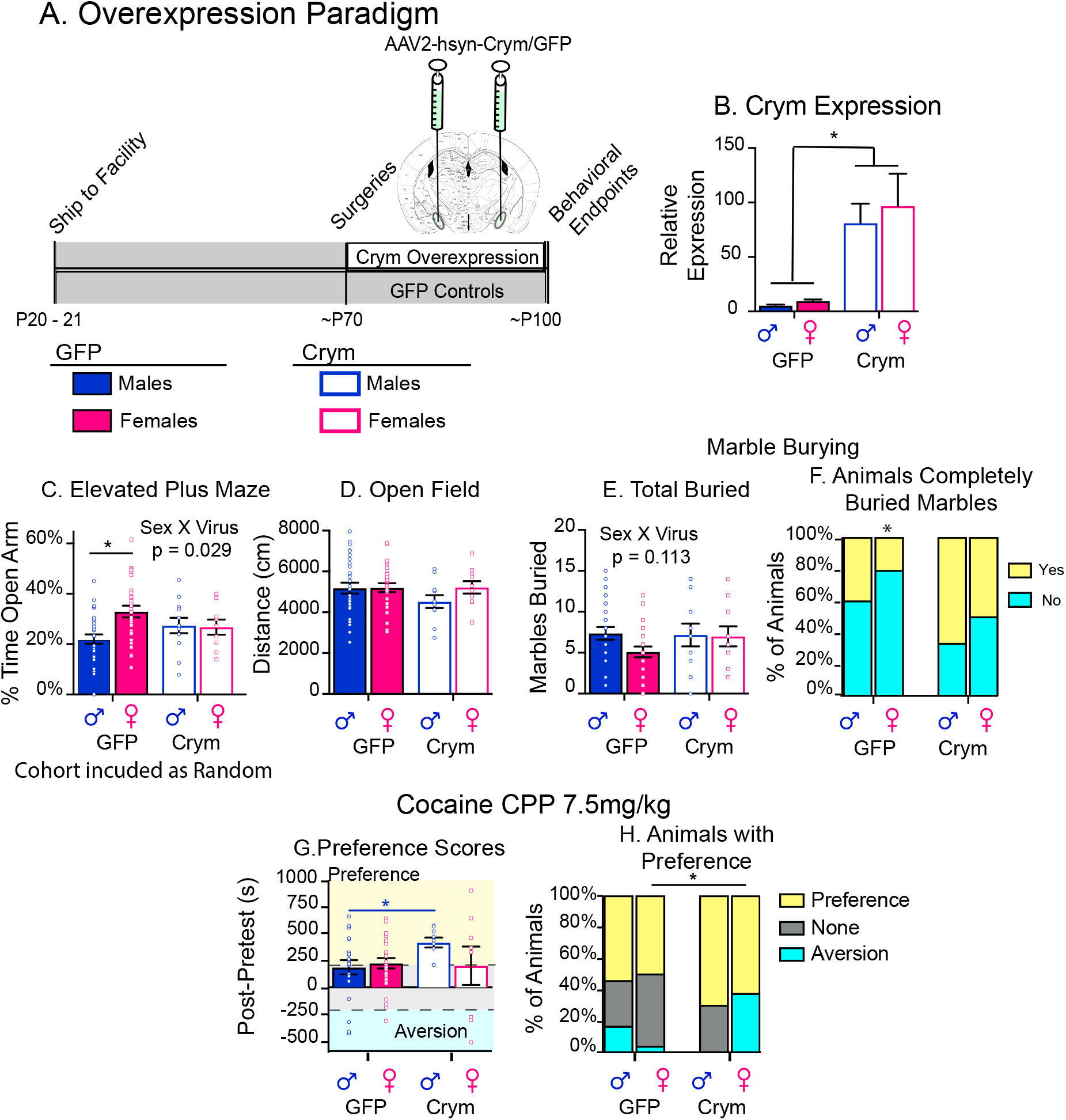
*Crym* OE in Adult meA Recapitulates Behavioral Effects of Adolescent SI. (A) Schematic of experimental design. (B) Viral expression and placement were confirmed after behavioral testing by epiflourescence microscopy (not shown) and qPCR of meA. (C-F) Similar to adolescent SI, *Crym* OE in meA abolishes known sex differences in elevated plus maze (C) and marble burying (E-F), but has no effect on distance traveled in an open field (D). (G-H) Similar to adolescent SI, *Crym* OE in meA increases cocaine CPP in males. While *Crym* does not replicate the effects of SI in females (G), it does increase the number of females who form an aversion to cocaine (H, blue). Additionally, *Crym* OE results in a bimodal distribution in cocaine CPP in females, which display either a preference (yellow) or aversion (blue) for cocaine. Post-hoc significant effects indicated as *p<0.05; **p<0.001. See also Suppl Figure 6.

Since SIM displayed increased *Crym* expression in meA after acute cocaine (Figure 5F), we hypothesized that *Crym* OE would recapitulate the SI-induced increase in cocaine CPP in males. As predicted, there were no differences in cocaine CPP between GFPM and GFPF, and *Crym* OE increased cocaine CPP in males compared to GFP controls (p=0.04). However, *Crym* OE in females caused an unexpected bimodal distribution for cocaine CPP: females either formed a preference for cocaine or an aversion (Figure 6G,H). Because of this highly variable bimodal distribution, non-parametric testing was applied and males and females were analyzed separately. Analysis of the distributions revealed a significant difference across the groups (Pearson’s Chi Squared: χ^2^=13.4; p=0.05), and analysis between groups revealed a significant difference between GFPF and CrymF (p<0.01) but not CrymM vs. F (p= 0.08) (Figure 6H). Importantly, when we overexpressed another gene of interest, *Lhx8*, in meA that does *not* meet criteria for a sex-specific key-driver gene (Suppl Figure 5A-H), sex differences are maintained in EPM (Suppl Figure 6B) and marble burying (Suppl Figure 6D,E) and no effects on cocaine CPP are observed (Suppl Figure 6E,F). These data thereby validate our network analysis and criteria for determining sex-specific key driver genes.

To determine if *Crym* action is specific to meA, we overexpressed *Crym* in nucleus accumbens (Suppl Figure 6H) and found no effects on OF (Suppl Figure 6I), marble burying (Suppl Figure 6J,K) or cocaine CPP (Suppl Figure 6L,M). This suggests that the effect of Crym expression specifically in meA is responsible for regulating sex-specific anxiety- and reward-associated behaviors.

### *Crym* OE in adult meA mimics transcriptional effects of adolescent SI

We next sought to validate *Crym’s* role as a key - driver gene at the transcriptional level. RNA-seq was conducted on punches of meA in which *Crym* OE was validated by qPCR (Figure 6B). We first compared sex differences in DEGs in GFPM and GFPF to those in *Crym* OE mice and found that, as with SI, *Crym* OE induced loss of expected sexually-dimorphic expression (Figure 7A,B; Suppl Table 3) as well as a gain of sex differences (Figure 7B; Suppl Figure 7A). Union heatmaps of DEGs altered by *Crym* OE in males vs. females (compared to GFP controls) revealed that *Crym* OE results in more DEGs in males than females (Figure 7C), with very little overlap between the two sexes (Figure 7D). These data are in line with transcriptional effects observed after SI (Suppl Figure 2).

**Figure 7:**
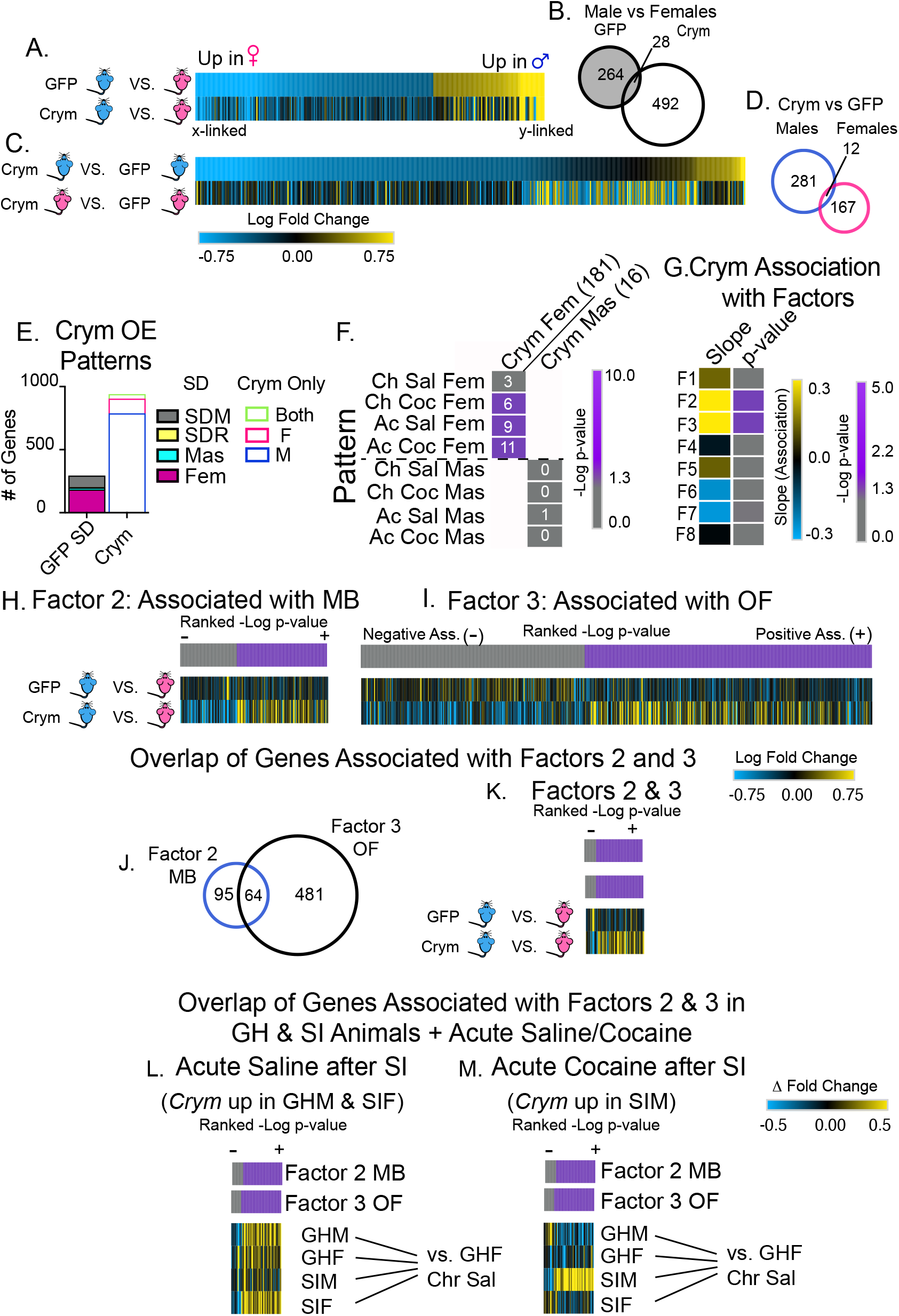
*Crym* OE in Adult meA Recapitulates Transcriptional Effects of Adolescent SI. (A-D) Heatmap of sexually-dimorphic genes in GFP animals (A). *Crym* OE abolishes expected sex differences, but induces new ones (B). (C-D) Union heatmaps and Venn diagrams of all DEGs in males and females after *Crym* OE. As with SI, *Crym* OE affects more transcripts in males than females and there is very little overlap of DEGs between the sexes. (E) Pattern analysis of *Crym* transcriptional effects. DEGs compared to the same baseline (GFPF) and categorized as previously discussed. Number of sexually-dimorphic genes categorized as each SD pattern (left) or genes only affected by Crym OE categorized as each pattern. (F) Enrichment of genes categorized as feminized or masculinized after adolescent SI or Crym OE. Grids show pairwise comparisons of gene lists. Purple = significant enrichment of the two lists (FDR p<0.05). Number in each box = number of overlapping transcripts. Numbers in parentheses = total number of genes. (G) Factors correlated with behavioral endpoints (F1-F8) and their associations with Crym OE in meA. Positive = yellow; negative = blue; significant association = purple (G). (H-I) Heatmaps of genes significantly associated (p<0.05, slope 0.15) with Factors 2 (H) or 3 (I) and their expression patterns in GFPM vs. GFPF or CrymM vs. CrymF. Genes are organized by the ranked −log p-value of the positive (purple) or negative (gray) association with each Factor. Associations with each factor are driven by sex differences in expression between CrymM vs. CrymF. (J) Venn diagram of the overlap of genes associated with Factors 2 or 3. (K) Heatmap of the 64 genes that overlap between Factors 2 and 3 and their expression in GFPM vs. GFPF and CrymM vs. CrymF. Genes are organized by their ranked −log p-value of positive (purple) and negative (gray) associations with Factor 2. (J-M) Heatmaps of the 64 genes that are associated with both Factors 2 and 3 and their expression in GHM/F or SIM/F after acute saline (J) or acute cocaine (M). Genes are organized by their ranked −log p-value of positive (purple) and negative (gray) associations with Factor 2. Associations are reflected in expression profiles in GHM and SIF after acute saline and SIM after acute cocaine. See also Suppl Figure 7 and Suppl Tables 3 and 4.

We conducted pattern analysis to determine if *Crym* OE feminized or masculinized sexually-dimorphic expression or induced sex - specific effects on transcription. Similar to the patterns observed after SI, we found that a large proportion of sexually-dimorphic genes were feminized by *Crym* OE and that the vast majority of gene changes induced by *Crym* OE occur in males (Figure 7E; Suppl Table 3). Enrichment analysis revealed significant overlap of genes categorized as feminized by adolescent SI under all treatment paradigms except chronic saline and those categorized as feminized by *Crym* OE, but no enrichment of genes categorized as masculinized (Figure 7F). These data suggest that *Crym* OE recapitulates the transcriptional profiles in SI males, especially in response to acute stimuli.

Since *Crym* overexpression in meA induced similar effects on both behavior and transcription as SI, we tested the hypothesis that a suite of *Crym*-regulated genes are associated with sex-specific behaviors and that these genes respond to cocaine in a sex-specific manner. To test this, we employed a dimensional reduction technique, exploratory factors analysis, used previously by our laboratory to reduce the behavioral endpoints to factors which can then be compared to transcriptomic data to identify genes associated with each behavioral factor (Walker et al., 2018). This analysis revealed 8 factors associated with the behaviors, and factor loading (Figure S7A) suggested that Factors 1-4 were most strongly associated with key aspects of each behavioral test: Factor 1 is positively associated with time in open arm and negatively associated with time in closed arm of the EPM; Factor 2 is positively associated with all measures in marble burying; Factor 3 is positively associated with time in center and negatively associated with time in periphery of the OF; and Factor 4 is positively associated with cocaine CPP and negatively associated with pre-test values (Suppl Figure 7).

We next applied linear modeling to the RNA-seq data from GFP and *Crym* OE animals to identify transcripts that are positively or negatively associated with each Factor (Suppl Table 4). We focused on those factors with associations where *Crym* OE was significantly (slope<0.15; p<0.05) associated with the Factor (Figure 7G). *Crym* OE was positively associated with Factors 2 and 3 only (Figure 7G; Suppl Figure 7) – marble burying and open field, respectively – the two behaviors that were only subtly altered by *Crym* OE. The association of *Crym* OE with Factor 2 likely reflects that a greater proportion of *Crym* OE animals buried more marbles than GFP controls (Figure 6F), and the association of *Crym* OE with Factor 3 likely reflects subtle sex differences in OF behavior. We examined expression profiles of the transcripts positively (purple) and negatively (gray) associated with Factors 2 and 3 and noted that sex differences in *Crym* OE but not GFP mice were driving the associations with both behaviors (Figure 7 H,I). We also noted a high degree of overlap between the transcripts associated with both factors (Figure 7J). Therefore we hypothesized that the overlapping genes might reflect a suite of *Crym*-sensitive transcripts within meA that are responsible for driving sex differences in anxiety- and reward-associated behaviors (Figure 7K). Once again, genes overlapping between Factors 2 and 3 showed expression patterns that reflected sex differences in *Crym* but not GFP mice (Figure 7K).

Because *Crym* is upregulated by acute stimuli (saline or cocaine) in a sex-specific manner (Figure 5F), we hypothesized that these *Crym*-sensitive genes would be regulated in a sex-specific manner by acute saline or cocaine. We compared expression profiles of the *Crym*-sensitive genes in our initial RNA-seq data after acute saline or cocaine and found that their expression patterns reflected the positive and negative Factor associations in GHM and SIF after acute saline (Figure 7L) but not acute cocaine. In contrast, associations with acute cocaine were reflected in SIM expression (Figure 7M). Together, these data suggest that a suite of *Crym-*regulated transcripts in meA are important mediators of sex-specific anxiety- and cocaine-regulated behaviors and the sex-specific transcriptional response to cocaine.

Three genes associated with both Factors 2 and 3 (Suppl Table 4) piqued our interest because of their known function in reward and their sensitivity to adolescent stress: oxytocin (*Oxt*), vasopressin (*Avp*) and dopamine receptor 1 (*Drd1*). It is not surprising that these transcripts displayed sex-specific regulation by acute saline and cocaine as they are all important in reward (Holder et al., 2015; Keverne and Curley, 2004; Miller et al., 2019). However, the finding that *Crym* is a sex-specific regulator of all three is novel and deserves follow-up. Because *Crym* is a marker of activated *Drd1* neurons in nucleus accumbens after morphine (Avey et al., 2018), we hypothesized that transcripts associated with Factor 4, which is positively associated with cocaine CPP (Suppl Figure 7A), may be responsive to cocaine in a sex-specific manner. Indeed, this is what we found (Suppl Figure 7G,H). Many of the genes associated with Factor 4 are immediate early genes (Suppl Table 4) and have been implicated in regulating responses of *Drd1* neurons in nucleus accumbens to cocaine (*cFos, Fosb, Ppp1r1b, Arc*; (Mews et al., 2018). Together, these data suggest that *Crym* OE in meA reprograms expression of reward-associated transcripts, which leads to sex-differences in responses to cocaine.

## DISCUSSION

Here we integrated an adolescent manipulation with long-term behavioral and transcriptional effects, dimensional reduction bioinformatic techniques (Walker et al., 2018) and co-expression network analysis to identify a sex-specific key driver of anxiety- and cocaine-related behaviors in meA. In doing so, we highlight that adolescence is a sensitive window for development of key sex differences in anxiety- and reward-associated behaviors. We also characterize meA as a novel brain region to regulate sex-specific cocaine-related behaviors and show that sex differences in the transcriptome after SI reflect sex differences in behavioral effects of SI. We then establish that the transcriptional changes induced in meA by adolescent SI are meaningful for sex-specific behavior by targeting a key driver derived from our network analysis, *Crym*, specifically within me A and thereby recapitulating the behavioral and transcriptional effects of SI. Finally, we demonstrate that, by targeting a bioinformatically-deduced key driver in the adult animal, we can open a window of plasticity for sex-specific behaviors, and identified a suite of *Crym*-sensitive genes that may be critical for regulating such behaviors. This work supports the view that it may be possible to treat the lasting sequelae of adolescent trauma long after the experience itself, and highlights the power of an integrative approach for studying sex-specific effects of adolescence.

### Adolescent SI is a model for investigating disruption of sex-specific behaviors

Adolescent SI has been used for decades to influence anxiety- and drug-related responses, but very few studies have included females and most have not resocialized animals before testing. Since SI itself is stressful, not rehousing animals before testing makes it difficult to determine if its long-term effects are due to the stress of SI itself or if the adolescent period is a sensitive window for life-long effects (Walker et al., 2019). The present study is unique in its investigation of adolescence as a sensitive window for the development of sex-specific anxiety- and cocaine-associated behaviors in adulthood. Our findings show that adolescent SI has opposite effects in males and females in both behavioral domains.

We then leveraged the sex-specific behavioral effects of adolescent SI to investigate underlying transcriptional mechanisms. We focused on meA because of its known sex differences in size (Hines et al., 1992), transcription (Chen et al., 2019), development (De Lorme et al., 2012), mechanisms of sexual differentiation (Argue et al., 2017; Krebs-Kraft et al., 2010; Nugent et al., 2009; Zehr et al., 2006), importance in sex-specific reward-associated behaviors (Chen et al., 2019; Li et al., 2017; Unger et al., 2015) and sensitivity to adolescent stress (Cooke et al., 2000; Hodges et al., 2019). Additionally, while evidence suggests that meA is an important regulator of drug sensitivity in females (Rudzinskas et al., 2019), very little is known about males (Knapska et al., 2007). Therefore, we sought to broaden our understanding of baseline sex-specific transcriptional responses in meA to cocaine in addition to characterizing how those responses are affected by adolescent SI. In line with previous results investigating sex-specific transcription in other disease states (Barko et al., 2019; Hodes et al., 2015; Labonte et al., 2017; Pena et al., 2019; Seney et al., 2018), there is very little overlap in transcripts altered by cocaine in males vs. females, suggesting that control males and females respond very differently to the drug. Our pattern analysis reveals that these differences are not specific to cocaine, as robust sex differences in DEGs are seen after saline, presumably reflecting a stress effect.

Importantly, as reflected in Venn diagrams (Figure 2) and pattern analysis (Figure 3; Suppl Figure 3), these sex differences are lost after SI and new sex differences are gained. These SI-induced transcriptional changes thus parallel the behavioral changes induced by SI.

### *Crym* OE in meA recapitulates the behavioral and transcriptional profile of adolescent SI

We go on to show that the observed transcriptional changes induced by SI within meA are important for driving sex-specific behavior by manipulating a sex-specific key driver derived from our integrated bioinformatic techniques. We identified two potential key drivers, *Crym* and *Ptgds*, that met our criteria (Figure 5) and validated our bioinformatics approach by viral OE of *Crym*, which we selected based on its biological function as a non-canonical transcriptional regulator (Hallen et al., 2011). We show that *Crym* OE in meA of adult males and females recapitulates the effects of adolescent SI on behavior (Figure 6). By contrast, <u>adult</u> SI did not induce long-term changes in behavior (Suupl Figure 1), which suggests that targeting a key driver gene identified bioinformatically re-opened a window of plasticity in sex-specific behavior. We show that these effects are specific to meA as *Crym* OE in nucleus accumbens has no effect on behavior. Conversely, overexpression of *Lhx8*, a key driver that does not meet our criteria for driving sex-specific effects after adolescent SI, in meA had no effect on behavior (Suppl Figure 6).

*Crym* is an intercellular thyroid hormone-binding protein that prevents thyroid hormone from interacting with its receptors (Hallen et al., 2011). Thyroid hormone receptors are members of the nuclear receptor superfamily of transcription factors. When bound to DNA without ligand, thyroid hormone receptors are transcriptional repressors (Lazcano et al., 2019). Thyroid hormone has recently been proposed as a potential regulator of sex-specific critical windows of neuronal development (e.g., in the visual system) (Batista and Hensch, 2019). Our data support this hypothesis and expand on it by showing that *Crym* OE not only influences sex-specific behaviors but recapitulates the transcriptional changes induced by adolescent SI in meA as well (Figure 7). These data give credence to the hypothesis that thyroid hormone is indeed critical for the regulation of developmental windows and that, by disrupting such signaling in a sensitive brain region in adulthood, we can reopen that window. These observations highlight the enormous potential of our dataset in driving future research.

Finally, using exploratory factor analysis of behavior coupled with our RNA-seq data, we identified a suite of ~60 genes that may be important for the sex-specific behavioral effects of *Crym* (Figure 7; Suppl Figure 7). We also show that these genes display a sex-specific response to an acute saline injection and the opposite transcriptional response to cocaine in SI males. These data suggest that a small suite of *Crym*-responsive genes, possibly through thyroid hormone signaling, may be programmed during adolescence to drive sex-specific reward- and anxiety-related behaviors, programming that is disrupted by SI. While some of these genes have already been implicated in such behaviors, e.g., *Oxt*, *Avp*, and *Drd1*, our list also includes many novel transcripts that deserve follow-up. We then expanded on this finding by showing that those genes identified in our factor analysis as being positively or negatively associated with cocaine CPP are regulated by an acute injection of cocaine in a sex-specific way and that this association is disrupted by adolescent SI. Importantly, there is a larger number of predicted dopamine-responsive immediate early genes associated with Factor 4, including genes that encode DARPP-32 (PPP1R1B), cFos, and FosB (Robison and Nestler, 2011). Many of these genes are downregulated in meA by cocaine in GHF and SIM, and upregulated by cocaine in SIF. This may reflect an important sex-specific difference in second messenger signaling in response to dopamine within meA, as has been reported (Rudzinskas et al., 2019).

## Conclusions

Experience during adolescence programs meA to shape sex-specific behaviors that maximize reproductive success in adult individuals. While it is tempting to view anxiety- and reward-associated behaviors through the lens of psychiatric disorders, sex differences in these behaviors more than likely reflect differences in behavioral strategies used by males and females to find a mate. Although more research is needed, our data suggest that adolescent SI disrupts such behaviors through programming the meA transcriptome. Because adolescent animals integrate information about the environment to fine tune behaviors associated with reproductive strategies, we hypothesize that SI likely signals information about mate availability, social hierarchy, resource availability and the likelihood of communal nesting, all of which would be influenced by metabolic factors. Therefore, it is not surprising that *Crym*, a thyroid hormone-binding protein, would be important for reprogramming transcriptional responses for metabolically costly events (e.g., copulation). We hypothesize that *Crym* may be programming the sex-specific transcriptional response to acute stimuli in meA as a mechanism for driving sex-specific behaviors necessary for reproductive success. However, it is important to point out that this programming may induce changes that result in sex-specific susceptibility or resilience to psychiatric disorders including addiction and anxiety when environmental mismatches take place in adulthood. Our data thereby not only provide valuable insight into sex-specific transcriptional regulation of behaviors but also provide a model for studying sex-dependent and -independent mechanisms of vulnerability to psychiatric disorders.

## ACKNOWLEDGEMENTS AND FINANCIAL DISCLOSURES

This work was funded by grants from the National Institute on Drug Abuse (NIDA) P01DA0047233 (EJN), R01DA007359 (EJN), R01MH051399 (EJN) and K99DA042100 (NIDA). The authors reported no biomedical financial interests or potential conflicts of interest. We would like to thank Dr. Erin Kendall Braun for their thoughtful comments in editing the manuscript and Dr. Andrew Wolfe for technical and intellectual support in the experimental design.

## AUTHOR CONTRIBUTION

Conceptualization, D.M.W., B.Z., and E.J.N; Methodology, D.M.W, X.Z, A.R., H.M.C., G.E.H, L.S.; Investigation, D.M.W, H.M.C, A.M.C, C.J.P, R.C.B, O.I, Y.V, A.P.L, A.G., C.J.B, E.M.P, and A.T-B; Writing – Original Draft, D.M.W and E.J.N; Writing – Review & Editing, D.M.W, H.M.C, C.J.P. and E.J.N.; Funding Acquisition, D.M.W, B.Z. and E.J.N.; Resources, P.J.K, B.Z. and E.J.N.; Supervision, L.S., B.Z., and E.J.N.

## Methods

### Animals

All experiments were conducted in accordance with guidelines of the Institutional Animal Care and Use Committee at Mount Sinai. Age-matched male and female C57BL/6J mice were shipped from Jackson Laboratories (Bar Harbor, ME) and delivered to the facility on postnatal day (P) 21 or ~P60 (adult isolation controls). After 24 hr of acclimation, animals were isolated on P22 or ~P61 by placing one animal in a transparent polycarbonate cage. Animals had olfactory, visual, and auditory interactions with others but were not allowed tactile interactions. After three weeks of isolation (P42 or ~P81), animals were rehoused with their original cage mates and group housed (GH) until ~P90 or ~P120 when behavioral and molecular testing was conducted. For behavioral endpoints, 6 different cohorts of animals were used (3 cohorts for anxiety-related behaviors; 3 cohorts for cocaine CPP (n = 10-15/group/cohort)). For molecular endpoints, all animals were collected in one cohort. For each cohort, GH controls were maintained as comparisons. Animals were maintained on a 12 hr light-dark cycle (lights on at 7:00) at 22-25°C with *ad libitum* access to food and water.

#### Vaginal Cytology

For all endpoints, vaginal cytology was monitored prior to and during behavioral testing and was included in the analysis to confirm that estrus cycle had no effect on the above behavioral endpoints.

### Behavioral Testing

#### Elevated Plus Maze (EPM)

Sex differences in EPM are well established (Archer, 1975; Goel and Bale, 2009; Voikar et al., 2001) and have been used previously as a measure of hormonal reorganization of sexually dimorphic behaviors (Palanza et al., 2008). Generally speaking, females spend more time in the open arm of an EPM (Archer, 1975; Goel and Bale, 2009; Voikar et al., 2001). On the day of testing, animals were moved into a separate testing room and given at least 1 hr to habituate before testing commenced. Males and females were run on 2 different acrylic EPMs standing ~92 cm tall with 4 cross shaped arms (12 × 50 cm), 2 arms enclosed with opaque walls (40 cm tall) and 2 open arms. Males were run on one maze and females were run on the other under dim red light. Animals were placed at the center of the maze facing one of the open arms and behavior was recorded for 6 min and tracked using Ethovision 13.0. Amount of cumulative time spent in the open arms, closed arms, center, and total time tracked were recorded. In a few cases during testing, the mouse was not detected by the tracking software because of a shadow from the closed arm. Therefore, the total tracking time was used to calculate percent time in the open and closed arms. If tracking was lost for greater than 20% of the total time, the data were not used in the analysis. In the experiment examining Crym OE in the NAc, tracking was lost (>20%) in over half of the animals. Therefore, this test was excluded from analysis. All testing was conducted between 10:00 and 17:00 hr and groups were balanced throughout the testing.

#### Open Field testing (OF)

Sex differences in OF are sensitive to the time of day, species, and strain (Swanson, 1969; Voikar et al., 2001) but as with EPM, this behavioral test has been used as a measure of disruption of sex-specific programming of behavior by hormones (Goel and Bale, 2008; Rubin et al., 2006). On the day of OF testing, animals were moved into a separate testing room and given at least 1 hr to habituate before testing commenced. Animals were placed in the corner of an arena (44 × 44 cm) and allowed to explore the open arena for 10 min. Behavior was recorded and tracked using Ethovision 13.0 and time in the center and periphery as well as distance and velocity travelled were recorded. Once again, males and females were run in separate arenas and groups were balanced throughout the testing day. All testing was conducted between 10:00 and 17:00 hr under dim red light.

#### Marble burying

Sex differences in marble burying are well established (Goel and Bale, 2008; Wilson et al., 2004) and sensitive to reorganization by gonadal hormones during puberty (Goel and Bale, 2008). Males bury more marbles in this task than females and testosterone exposure during puberty leads to an increase in burying in females (Goel and Bale 2008). On the day of testing, animals were moved into a separate testing room and given at least 1 hr to habituate before testing commenced. Animals were placed in a plexiglass cage (19.56 × 30.91 × 13.34 cm) containing 9 cm of corn cob bedding and 20 sterilized glass marbles (13 mm diameter) placed in a 4 × 5 grid and evenly distributed across the entire cage. Animals were placed in the center of the arena and a lid was placed gently on the cage. After 15 min, animals were removed from the testing arena and the number of marbles partially and completely buried were counted and recorded. Marbles were counted by two researchers blind to treatment group. Each animal was run in its own cage with clean bedding and marbles. All testing was conducted between 10:00 and 17:00 hr under dim red light and groups were balanced throughout the testing day.

#### Cocaine conditioned place preference (CPP)

Sex differences in cocaine-related behaviors, including CPP, are less consistent and appear to be dependent on dose, experimental procedure and species. However, it is notable that, when sex differences are observed, females form a greater preference for cocaine compared to males (Calipari et al., 2017; Russo et al., 2003; Zakharova et al., 2009). On the days of testing, animals were moved into a separate testing room and given at least 1 hr to habituate before testing commenced. Three chamber CPP apparati were used with 2 large outer chambers and one small middle chamber. All chambers differed in tactile and visual cues. One large chamber had striped walls and small wire mesh floors, while the other had gray walls and large wire mesh floors. The middle smaller chamber had white walls, metal rods as the floor and two overhead lights (vs. one in each of the larger chambers). During pretesting, animals were placed in the middle chamber and were allowed to explore all three chambers for 20 min. Infrared beam breaks were monitored using Med Associates (San Diego, CA) software to track locomotor activity and time spent in each chamber. After pretesting, pre-CPP score was calculated as time spent in the gray chamber minus time spent in the stripe chamber. Any animals spending greater than 300 sec on one side during the pre-test were excluded from the study, which amounted to <10% of all animals.

An unbiased approach was utilized to determine conditioning. For each group, an equal number of animals were conditioned with cocaine on their preferred or non-preferred side. On each of two conditioning days animals were injected with saline between 10:00-13:00 hr and placed in one chamber for 30 min. In the afternoon (14:00-18:00), animals were injected with cocaine (7.5 mg/kg) and placed in the opposite chamber for 30 min. The following day, animals were placed in the apparatus and allowed equal access to all three chambers. An animal was considered to have a preference for cocaine if they spent more than 50% (>625 sec) of their time in the cocaine-paired chamber or considered to have an aversion to cocaine if they spent more than 50% of their time during testing on the saline-paired side. Half of the males in each cohort were run in one week followed by females over the next ~10 days. The second half of the males were run after the females were completed to account for any time that passed during testing. Groups were balanced between testing conditions and time of testing was included in the analysis to confirm that time was not a factor in the development of CPP. For females, vaginal cytology was monitored 7-10 days before behavioral testing commenced to confirm that females were cycling regularly. The pretest was conducted on one day for all of the females. However, because evidence suggests that cocaine CPP is dependent on cycle stage in female mice (Calipari et al., 2017), females were conditioned when they were in proestrus and estrus and testing occurred ~24 hr after the last conditioning day. This means that females were in metestrus or diestrus I when behavioral testing occurred, during which time gonadal hormones were likely decreasing.

### Statistical Analysis of Behavioral Data

All behaviors were analyzed using a 2-way ANOVA or a Kruskall-Wallis non-parametric test depending on the significance of a Levene’s test for homogeneity of variance. If an interaction was identified, a Tukey post-hoc analysis was conducted to determine specific differences between groups. If an effect was identified via Kruskall-Wallis, follow-up Mann-Whitney tests were run to determine specific differences between groups. Chi-square tests were used to determine differences in the distribution of behavioral phenotypes within a group (marble burying and CPP). Pearson’s Chi Squared tests were used to account for the lack of representation of all categories across groups. All analyses were conducted using SPSS Statistical Software, V25 (IBM, Armonk, NY).

### Cocaine Injections and Tissue Collection for RNA-seq

On P80, animals were divided into 8 groups of males and 8 groups of females: GH + chronic cocaine/saline; SI + chronic cocaine /saline; GH + acute cocaine/saline; SI + acute cocaine/saline. In total, 200 animals were utlilized. For chronic injections, animals were given one injection (IP) of cocaine or saline per day for 10 days (between 10:00 and 14:00 hr) and were euthanized 24 hr after their last injections (n=6-8 animals/group; total = 116 samples). For acute injections, animals were given saline injections (IP) for 9 days in an effort to habituate the animals to handling and injection stress. On the 10^th^ day, animals were injected with saline or cocaine (7.5 mg/kg) and euthanized ~1 hr after the injection. All animals were euthanized via cervical dislocation. Brains were removed and sectioned on ice in a brain block (1 mm thick) and bilateral micropunches (15 gauge) of meA were snap frozen on dry ice and stored at −80°C until use. Vaginal cytology was monitored throughout the injections and only those females in metestrus/diestrus were used for library preparation.

### RNA Isolation, Library Preparation, and Sequencing

RNA was isolated as described previously (Walker et al., 2018) using RNAeasy Mini Kit (Qiagen, Fredrick, MD) with a modified protocol from the manufacturer allowing for the separation and purification of small RNAs from total RNA. Briefly, after cell lysis and extraction with QIAzol (Qiagen, Fredrick, MD), small RNAs were collected in the flow-through and purified using the RNeasy MinElute spin columns and total RNA was purified using RNeasy Mini spin columns. Samples were treated with DNAse to rid them of genomic DNA and run on nanodrop and an Agilent Bioanalyzer 2100 to confirm RNA purity, integrity and concentration. All samples exhibited RINs >8.

Libraries were prepared using the TruSeq Stranded mRNA HT Sample Prep Kit protocol (Illumina, San Diego, CA). Briefly, poly A selection and fragmentation of 300 ng of RNA was performed, and the resulting RNA was converted to cDNA with random hexamers. Adapters were ligated and samples were size-selected with AMPur XP beads (Beckman Coulter, Brea, CA). Barcode bases (6 bp) were introduced at one end of the adaptors during PCR amplification steps. Library size and concentration were assessed using Tape Station (Life Technologies, Grand Island, NY) before sequencing. Libraries were pooled for multiplexing (pool of 21 samples) and sequenced on a HighSeq2500 System using V4 chemistry with 50 base pair single-end reads at UCLA Neuroscience Genomics core. Each pool was sequenced 3 times with the goal of obtaining ~30 million reads per sample. QC revealed an average of 29 million reads per sample (min = 19 million; Max = 51 million) with and average mapping rate of 90.2%. The number of independent tissue samples included in the final analysis was between 6-8 per group.

### Differential Expression Analysis

Gene expression raw read counts were normalized as counts per million (CPM) using trimmed mean of M-values normalization (TMM) method (Robinson et al., 2010) to adjust for the differences in library size among samples. Genes expressed at ≤1 CPM in ≥5 samples were removed before further analysis. Differentially expressed genes (DEGs) in comparisons of male vs. female, SI vs. GH, cocaine vs. saline and acute vs. chronic were identified using the Bioconductor package *limma* (Ritchie et al., 2015) with the following thresholds: nominal significance of p<0.05 and fold-change of 30% (Suppl Table 1).

### Pattern Analysis

Pattern analysis was conducted as described previously (Walker et al., 2018). Briefly, expression changes from the same baseline were converted into 0s and 1/−1 (0 = no effect; −1 = significantly downregulated; 1 = significantly upregulated) for each condition. Output from the analysis resulted in a list of combinations of 0s and −1/1s observed in the dataset. The patterns of expression were defined by two investigators unfamiliar with the experiment to avoid bias. Each gene was then assigned to a defined category. Patterns included genes that were differentially expressed when compared to GHF injected with chronic saline (p<0.05; fold change >30%). While many patterns were observed in the dataset (Suppl Table 2), we focused on two: those genes that were differentially regulated between GHM vs. F under the each exposure paradigm (sex differences; Figure 3) and those that were only affected in SI animals (SI-only Effects; Figure S3). For example, a gene that was significantly increased in GHM but not GHF after acute saline would be categorized as a sex difference in expression; if that gene’s expression resembles GHF in SIM then it was categorized as “feminized”, whereas if that gene resembles the expression profile of GHM in SIF it was categorized as “masculinized.” A schematic of the different categories is included in Figure 3A-D (sex differences) and Suppl Figure S3 (SI-only). Importantly, a gene can only be defined as one category under each treatment condition. Thus, we identified genes that are uniquely regulated by each stimulus.

### Multiscale Embedded Gene Co-Expression Network Analysis (MEGENA)

MEGENA (Song and Zhang, 2015) was used to construct gene co-expression networks for GHM, GHF, SIM and SIF samples. First, MEGENA constructed a planar filtered network by utilizing parallel computation, early termination and prior quality control. Next, MEGENA performed a multiscale clustering analysis to obtain co-expression modules by introducing compactness of modular structures characterized by a resolution parameter. Lastly, MEGENA conducted a multiscale hub analysis to identify highly connected hubs (or driver genes) of each module at each scale.

DEGs identified in the differential expression analysis were then laid onto the modules to perform enrichment analysis: namely, ranking each modules associated with different treatment conditions according to the adjusted enrichment p-value. In addition, GO-function enrichment analysis (Wang et al., 2012) was applied to the modules to identify enriched biological processes. Sunburst plots showing the enrichment of sexually dimorphic (SD) changes, SI changes or GO-biological processes in individual modules of the networks were visualized using the R package *sunburstR*. Top-ranked modules in each network were also visualized in circos plots with the significance of DEG enrichment using R *Netweaver.*(Wang et al., 2016). Module subnetworks were visualized by Cytoscape_v3.3 (Shannon et al., 2003).

Module differential connectivity (MDC) (Zhang et al., 2013) analysis was used to quantify network module reorganization across different conditions. MDC is a measure of the ratio of the average connectivity among genes of a module in one condition to changes among the same set of genes in another condition.

### Enrichment Analysis

Fisher’s exact tests (FETs) were conducted using the Super Exact Test package in R as described previously to determine module preservation as well as enrichment of patterns and other gene lists (Walker et al., 2018; Wang et al., 2015).

### Exploratory Factor Analysis

Exploratory factor analysis was conducted as described (Walker et al., 2018). Briefly, factor analysis was used to reduce the dimensions of the interdependent behavioral variables and help account for variability in the data due to differences in cohorts and viral expression. All behavioral data were converted to non-negative values followed by log2(x+1) transformation. A standard factor analysis was performed using the scikit-learn package (Pedregosa et al., 2011).

A 10-fold cross-validation (CV) was utilized to choose the number of factors. We found that the CV log-likelihood was maximized with 8 factors. Therefore, the factor number was set to 8 and factor analysis was then applied to the whole dataset. The transformed data from the analysis was used as a continuous variable for each factor. Differential analysis was conducted using Voom Limma to determine which factors were associated with gene expression (Law et al., 2014). Transcripts were considered associated with a factor if the slope of the association was at least 0.15 and the nominal p-value for the association was p<0.05.

### Viral Vectors

*Crym* and *Lhx8* cDNA was synthesized and cloned into an pAAV plasmid (Gene Script). The plasmid was package into an AAV2 vector by the Duke viral vector core (Durham, NC). This vector expresses both the gene of interest and eGFPunder the *hSyn* promoter and were separated by a t2a. AAV2-hSyn expressing eGFP only was used as the control.

For *in vivo* behavioral validation of key driver genes, male and female C57BL/6J mice were injected with AAV2 vectors and, ~3 wks after surgery when transgene expression is maximal, mice were run through all behavioral paradigms discussed above. Briefly, animals were shipped from Jackson Laboratories at ~P21 and allowed one week to acclimate to the new facility. On ~P70 we overexpressed (OE) genes of interest via stereotaxic injections of AAV2s expressing Crym or Lhx8 plus GFP, or GFP alone, in meA under ketamine (100 mg/kg IP)-xylazine (10 mg/kg IP) anesthesia. Animals were placed in a small-animal stereotaxic instrument (Kopf Instruments, Los Angeles, CA). Vectors (0.5 µl of viral titer = 1*10^12 particles/ ml) were bilaterally injected using 33-gauge syringe needles (Hamilton) at a rate of 0.1 µl/min into meA (Bregma coordinates: anterior/posterior, −1.1; Medial/lateral, 2.5mm; dorsal ventral, −5.3mm; 0° angle) or NAc (bregma coordinates: anterior/posterior, 1.6; Medial/lateral, 1.5mm; dorsal ventral, −4.4mm; 10° angle). Injections were performed in three cohorts with viruses represented across each cohort. Behavioral testing was conducted as described above, however, to limit the number of animals all 4 behaviors were run in each animal in the following order: week 1: EPM, OF, marble burying; week 2-3: cocaine CPP. As stated above, males were run first in cocaine CPP and females were run the following week. All females were trained on proestrus and estrus and CPP testing was conducted when females were in metestrus/diestrus 1. A pilot experiment was run prior to surgeries in a separate cohort of animals to confirm that CPP behaviors were not affected by previous behavioral testing or handling. Behavioral endpoints were analyzed as described above. Cohort was included as a factor in the analysis. If a cohort effect was observed for an endpoint, the direction and effect was confirmed to be the same for each cohort. If the patterns for the behaviors were the same across cohorts but small differences were observed in behaviors, cohort was included as a random factor in the ANOVA. For the cohort of animals with *Crym* OE in NAc, CPP testing was conducted in a different room using different conditioning chambers. A small but significant difference in behavior was noted in these testing chambers. Therefore, a slightly different threshold was applied to the preference scores for this cohort only: preference = ≥50% time (600 sec) in the cocaine-paired chamber (rather than >50% time in the cocaine-paired chamber).

### RNA Isolation and qPCR Validation

Animals were euthanized by cervical dislocation ~10 days after the last day of behavioral testing. Brains were removed and sectioned in a brain matrix (1 mm sections) and viral vector expression was confirmed by epifluorescence microscope. Only those animals that displayed GFP expression in meA were processed for RNA isolation. RNA was isolated as described above and converted to cDNA using High Capacity Reverse Transcriptase Kits (Catalog #4368814; ThermoFisher, Foster City, CA) according to manufacturer’s protocol. qPCR was performed for 2 genes of interest and 2 internal controls (Suppl Figure S1) using Taqman**®** gene expression assays (Thermo Fisher, Foster City CA) and Taqman**®** Fast Universal Master Mix (ThermoFisher, Foster City, CA) on an ABI Quant Studio Flex 7 according to the manufacturer’s protocol and using the following parameters: 1 cycle (2 min @ 50°C followed by 2 min @ 95°C); 45 cycles (1 sec @ 95°C followed by 20 sec @ 60°C). Expression was analyzed for all three viral vectors in each animal using the comparative ΔCt method (3). Each sample was normalized to its own internal controls (geometric mean of the Ct values for *Hprt1* and *Actb)* and calibrated to the average ΔCt for the GFP groups. Only those animals that had expression values >1.5 SD from the mean expression of *Crym* or *Lhx8* in the GFP groups were used in the behavioral analysis and the follow-up RNA-seq.

### RNA-Seq Analysis for *Crym* OE

Aliquots of RNA with confirmed Crym OE (or GFP controls) were sent to GeneWiz (New Jersey) for library preparation and RNA-seq (total N = 68). Paired-end sequencing reads were aligned to mouse genome GRCm38.95 using STAR aligner (Dobin et al., 2013). FeatureCounts (Liao et al., 2014) was then used to assign reads to genes for quantifying gene expression levels based on GENCODE (Frankish et al., 2019) release M20. Read counts were normalized as CPM using TMM method to adjust for differences in library size among samples. Principal component analysis and unsupervised clustering analysis were applied to the normalized expression levels to assess whether there were any outliers. Genes expressed at ≥1 CPM in ≥5 samples were obtained for DEG analysis. Limma was applied to the normalized expression levels to identify DEGs between Crym OE and GFP in males and in females, as well as DEGs between male GFP vs. female GFP, and male *Crym* OE vs. female *Crym* OE.

