## Supplemental Figures for "Adolescent Social Isolation Reprograms the Medial Amygdala: Transcriptome and Sex Differences in Reward"

**SUPPLEMENTAL INFORMATION:**

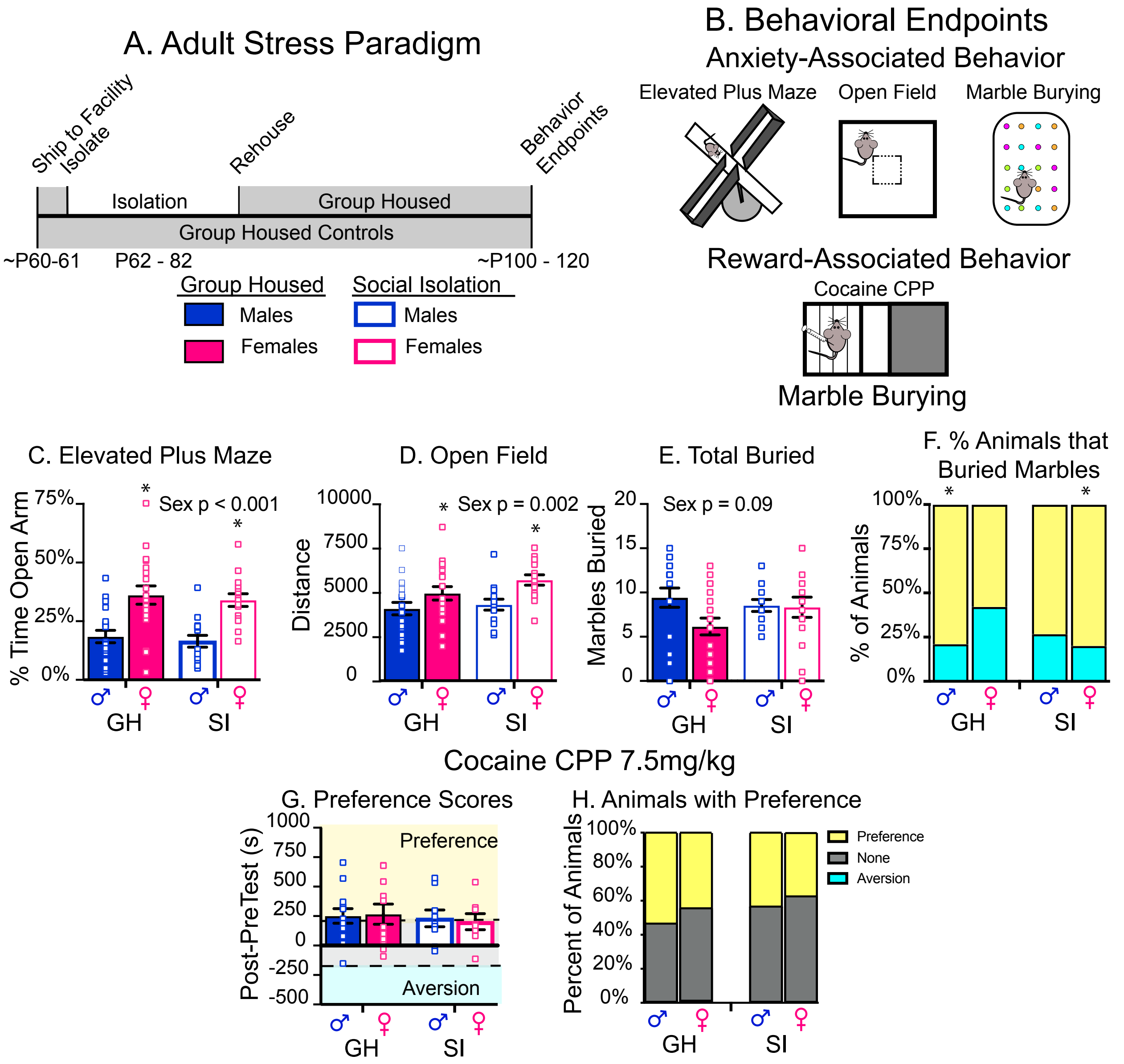


Figure 1: Adult Social Isolation Has No Effect on Sexually Dimorphic Behaviors or Cocaine CPP (A-B) Schematic of experimental design. (C – F) Adult SI results in a had no effects on sexually dimorphic behaviors in elevated plus maze (C) open field (D) and marble burying (E-F). (G-H) Adult SI has no effects on cocaine CPP (G) or in the proportion of animals that form a preference for cocaine in either males or females (H, blue).Post-hoc significant effects indicated as: * = p<0.05; ** = p<0.001.

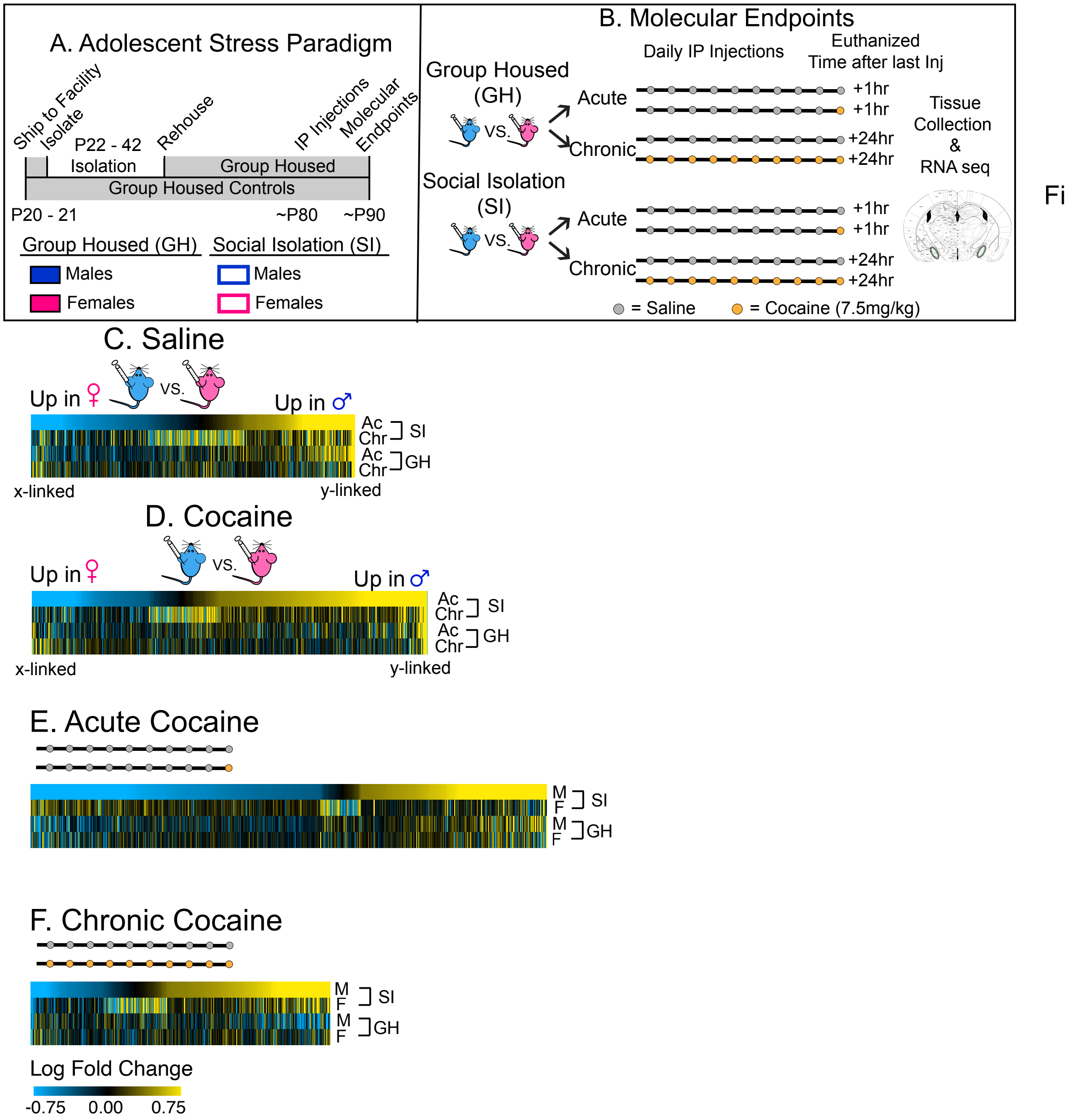


Figure 2: Adolescent SI Results in a SI specific transcriptional patterns in the meA.
(A-B) Schematic of experimental design. (C-F) Union heatmaps of all sexually-dimorphic genes significantly differentially expressed in SI males vs females after acute and chronic saline (C - D) and acute and chronic cocaine (E-F). There are more sex-differences in expression after an acute exposure to saline ( C & D) or cocaine (E & F) than under chronic conditions. Heatmaps reveal that adolescent SI results in the loss of sex-specific expression under all conditions.
(G-J) Union heatmaps of genes differentially expressed in SI males or females after acute (G-H) or chronic cocaine (I-J). Each comparison is compared to their own saline control. As indicated by the heatmaps, there is very little overlap of DEGs regulated by cocaine in SI males and females. Additionally, heatmaps reveal that SI results in an opposite transcriptional response to acute and chronic cocaine when compared to the GH males (G & H).

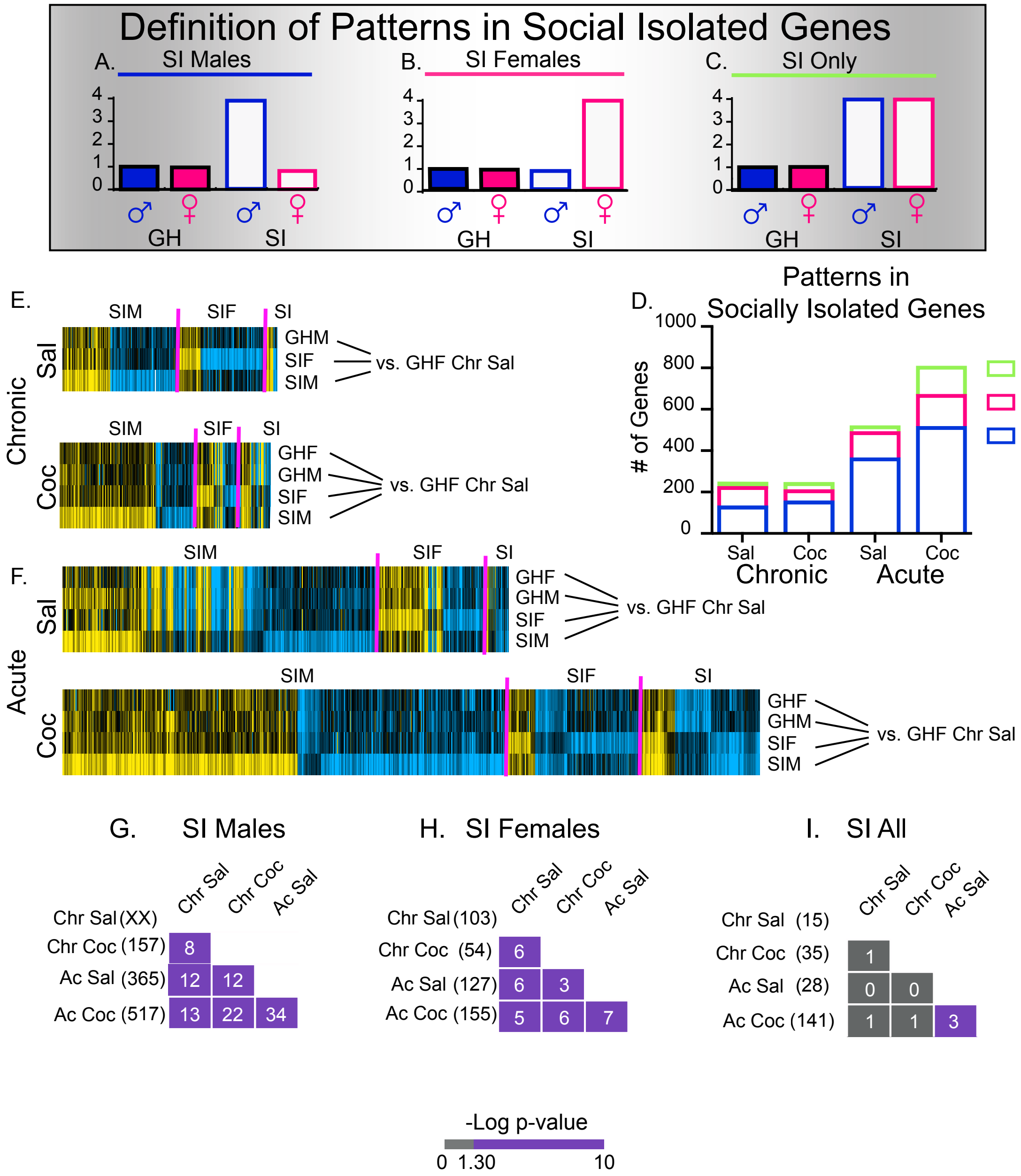


Figure 3: Pattern Analysis Reveals that Genes affected by SI.
(A-C) Theoretical Patterns: (A) SIM Only; (B) SIF Only; (C) SI Only; (E-H) Heatmaps of genes that fall into the categories of patterns represented in A – C. All comparisons are made to theGHF chronic saline expression and expressed as logFC. Pink lines indicate breaks the beginning of a new category; SIM = SI males only; SIF = SI Females only; SI = Both SI M & SIF but not GH animals. (I) Number of SI genes under each treatment paradigm and the number of transcripts that fall into each category. The number of SI genes increases with acute treatments of saline and cocaine in after SI. Across all treatment paradigms, the more genes were categorized as SIM. However after chronic saline, the percent of genes categorized as SIM and SIF are almost equal. (J) Enrichment plots showing the significant overlap (purple) of categories of genes across different treatment paradigms. Number of genes in each list are indicated in parentheses and the number of overlapping genes between conditions is indicated within the boxes. There is significant overlap of SIM and SIF genes across all treatment paradigms.

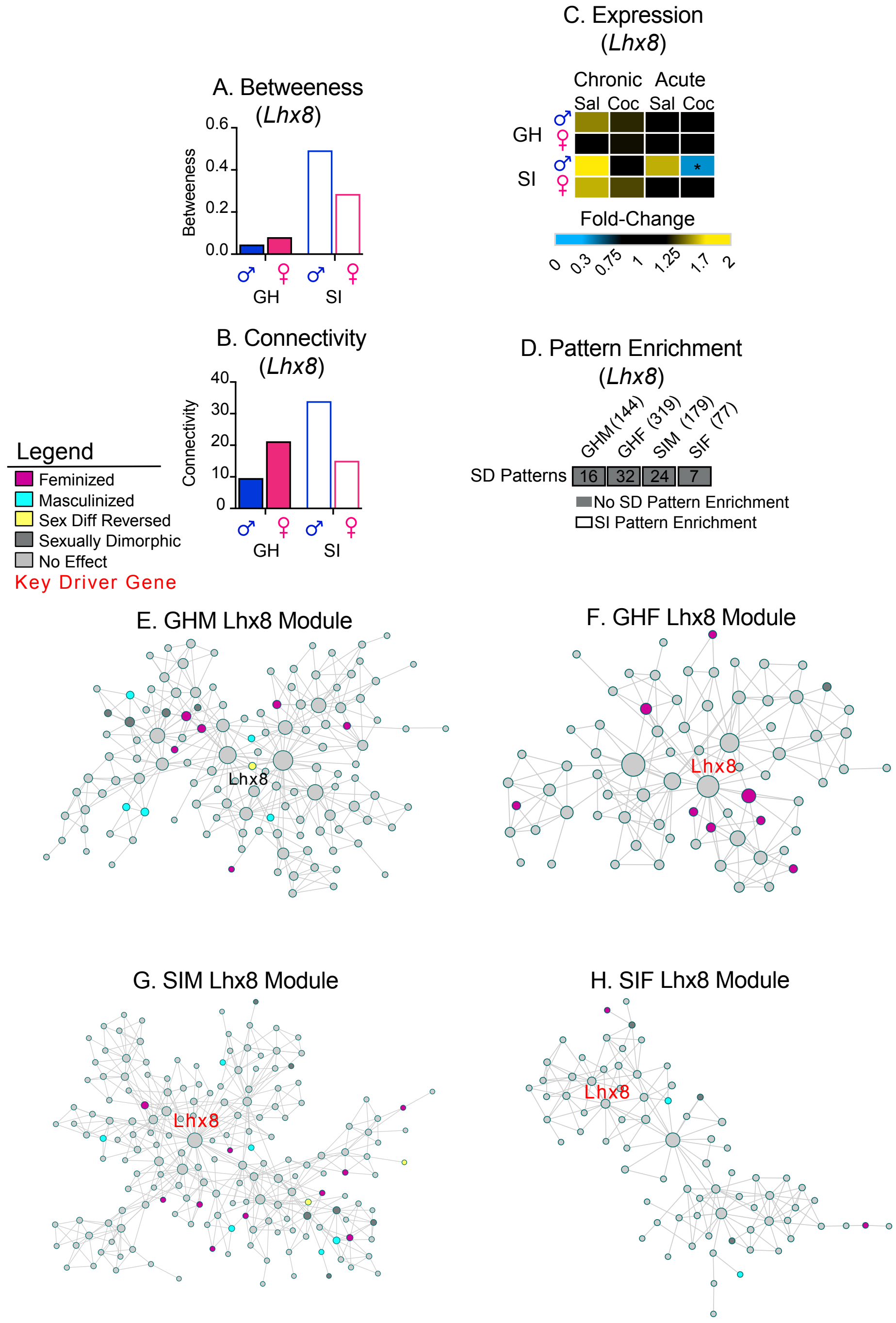


Figure 5: The transcription factor, *Lhx8*, does not meet the criteria used to identify sex-specific key drivers.

(A-C) LIM homeobox 8 (*Lhx8*) display increased betweenness (A) and connectivity (males only, B) after SI (open bars), and sex-specific expression patterns at baseline, which are lost after SI (C). Fold change in expression of *Lhx8* when compared to the same baseline (GHF after chronic sal). * indicate expression is significantly different from baseline and meets the 30% threshold for change in expression.(D) Enrichment of sexually dimorphic and SI only pattern genes is observed in the modules containing *Lhx8*. *Lhx8* containing modules are not enriched sexually dimorphic genes but are enriched in genes altered by SI males. (E-H) Arachne plots of *Lhx8* containing modules in all for groups. GHM (E), GH F (F), SIM (G) and SIF (H). Lhx8 is a key driver all modules except GHM as indicated by red text, sexually dimorphic genes falling into different patterns are color coded according to the patterns they display.

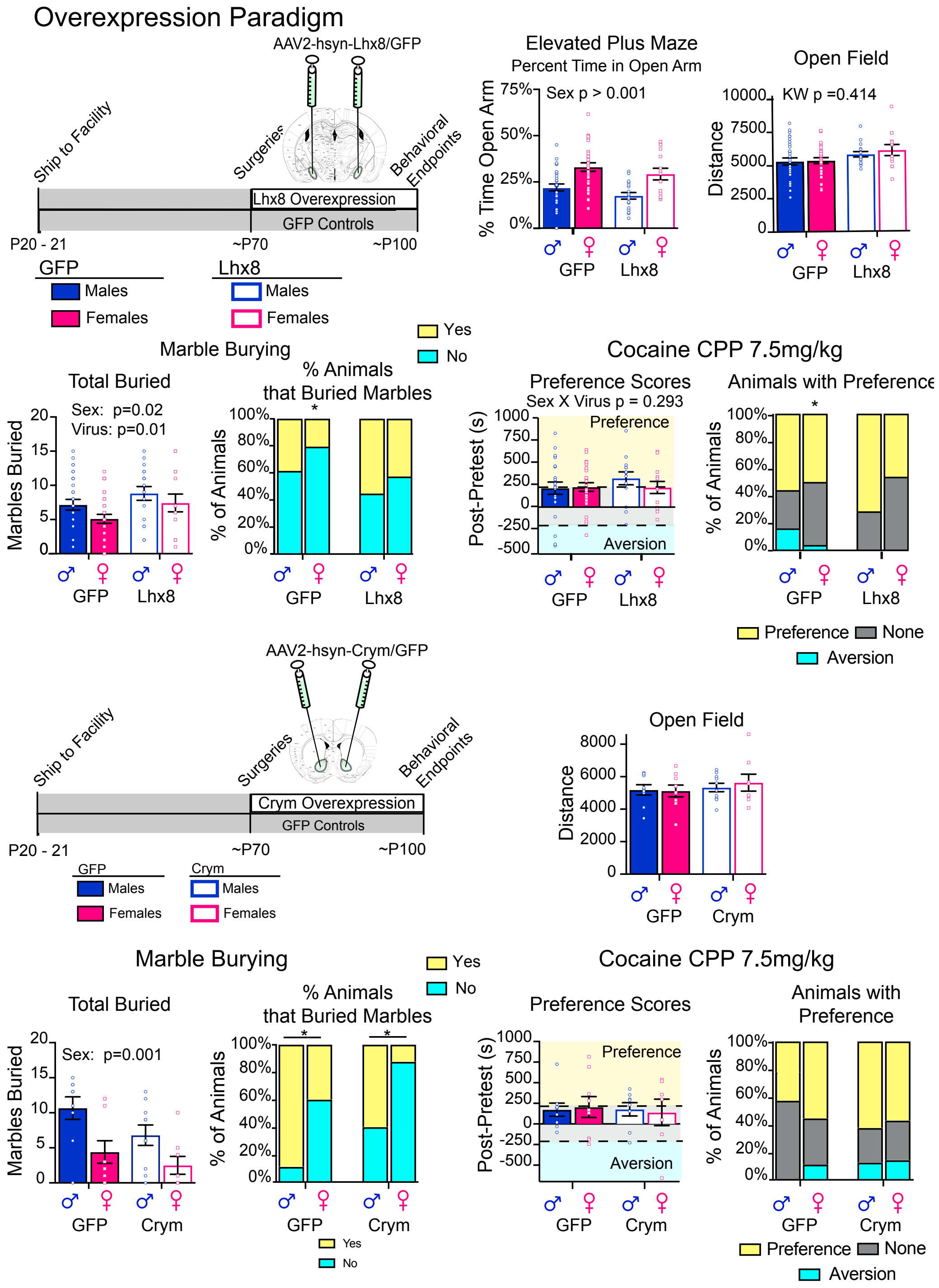


Figure 6 Lhx8 overexpression in the adult meA and *Crym* overexpression in the adult NAc has no effect on behavior. Schematic of experimental design for OE Lhx8 in meA. (B–G) *Lhx8* overexpression in the meA results has no effect on sex-specific behaviors in elevated plus maze (B) marble burying (D-E) and open field (C). It also has no effect on cocaine CPP (F-G) when compared to the GFP controls. However, *Lhx8* overexpression does result in an increase in the number of marbles buried in both males and females (D) and an increase in the proportion of animals that buried marble in males and females (E). (H) Schematic of experimental design for *Crym* OE in NAc. (I-M) *Crym* overexpression in the adult NAc has no effect on open field (I), marble burying (J-K), or cocaine CPP (L-M) when compared to GFP controls. Post-hoc significant effects indicated as: * = p<0.05; ** = p<0.001.

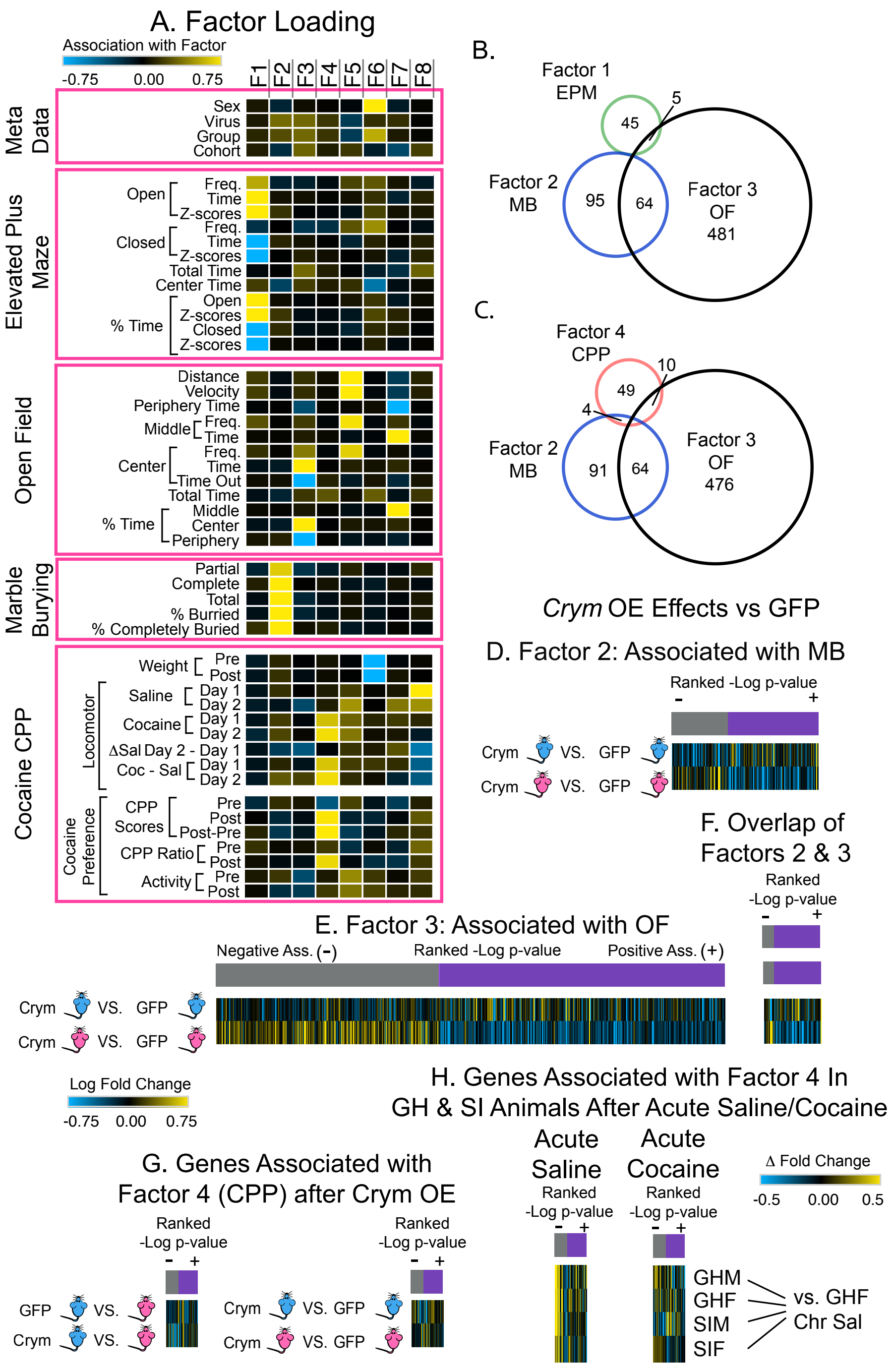


Figure 7: Factor Analysis Reveals Behavioral Endpoints Associated with *Crym* Regulated Genes.

(A) Factor loading of behaviors. Positive associations indicated in yellow; Negative associations indicated in blue; No association indicated in black. (B-C) Venn diagrams of transcripts associated with Factors 1 (EPM); 2 (MB) and 3 (OF) (B) or Factors 2; 3 and 4 (CPP). Heatmaps of genes significantly associated ($p<0.05$, slope 0.15) with Factors 2 (C) or 3 (D) and their expression patterns in CrymM vs. GFPM or CrymF vs. GFPF. Genes are organized by the ranked -log p-value of the positive (purple) or negative (gray) association with each Factor.

Figure 7 (Cont’d): (E) Heatmap of the 64 genes that overlap between Factors 2 and 3 and their expression in CrymM vs. GFPM and CrymF vs. GFPF. Genes are organized by their ranked -log p-value of positive (purple) and negative (gray) associations with Factor 2. (G) Heatmaps of genes significantly associated ($p<0.05$, slope 0.15) with Factors 4 and their expression patterns in GFPM vs. GFPF and CrymM vs. CrymF  or CrymM vs. GFPM and CrymF vs. GFPF. Genes are organized by their ranked -log p-value of positive (purple) and negative (gray). (H). Heatmaps of the genes associated with both Factors 4 and their expression in GHM/F or SIM/F after acute saline or acute cocaine. Genes are organized by their ranked -log p-value of positive (purple) and negative (gray). Associations are reflected in expression profiles in SIM after acute cocaine but not acute saline. Associations are not reflected in expression profiles in SIF which may provide insight into the bimodal distribution observed in behavior after Crym OE.
